# Temporal specificity of abnormal neural oscillations during phonatory events in Laryngeal Dystonia

**DOI:** 10.1101/2021.04.08.438908

**Authors:** Hardik Kothare, Sarah Schneider, Danielle Mizuiri, Leighton Hinkley, Abhishek Bhutada, Kamalini Ranasinghe, Susanne Honma, Coleman Garrett, David Klein, Molly Naunheim, Katherine Yung, Steven Cheung, Clark Rosen, Mark Courey, Srikantan Nagarajan, John Houde

**Author notes:** Correspondence to: Hardik Kothare or Srikantan Nagarajan or John Houde 513 Parnassus Ave, HSE 608, San Francisco, CA 94143, USA, Email addresses, and.

## Abstract

Laryngeal Dystonia is a debilitating disorder of voicing in which the laryngeal muscles are intermittently in spasm resulting in involuntary interruptions during speech. The central pathophysiology of laryngeal dystonia, underlying computational impairments in vocal motor control, remains poorly understood. Although prior imaging studies have found aberrant activity in the central nervous system during phonation in patients with laryngeal dystonia, it is not known at what timepoints during phonation these abnormalities emerge and what function may be impaired. To investigate this question, we recruited 22 adductor laryngeal dystonia patients (15 female, age range = 28.83-72.46 years) and 18 controls (8 female, age range = 27.40-71.34 years). We leveraged the fine temporal resolution of magnetoencephalography to monitor neural activity around glottal movement onset, subsequent voice onset and after the onset of pitch feedback perturbations. We examined event-related beta-band (12-30 Hz) and high-gamma band (65-150 Hz) neural oscillations. Prior to glottal movement onset, we observed abnormal frontoparietal motor preparatory activity. After glottal movement onset, we observed abnormal activity in somatosensory cortex persisting through voice onset. Prior to voice onset and continuing after, we also observed abnormal activity in the auditory cortex and the cerebellum. After pitch feedback perturbation onset, we observed no differences between controls and patients in their behavioural responses to the perturbation. But in patients, we did find abnormal activity in brain regions thought to be involved in the auditory feedback control of vocal pitch (premotor, motor, somatosensory and auditory cortices). Our study results confirm the abnormal processing of somatosensory feedback that has been seen in other studies. However, there were several remarkable findings in our study. First, patients have impaired vocal motor activity even before glottal movement onset, suggesting abnormal movement preparation. These results are significant because: (i) they occur before movement onset, abnormalities in patients cannot be ascribed to deficits in vocal performance, and (ii) they show that neural abnormalities in laryngeal dystonia are more than just abnormal responses to sensory feedback during phonation as has been hypothesised in some previous studies. Second, abnormal auditory cortical activity in patients begins even before voice onset, suggesting abnormalities in setting up auditory predictions before the arrival of auditory feedback at voice onset. Generally, activation abnormalities identified in key brain regions within the speech motor network around various phonation events not only provide temporal specificity to neuroimaging phenotypes in laryngeal dystonia but also may serve as potential therapeutic targets for neuromodulation.

## Introduction

Laryngeal Dystonia (LD), or Spasmodic Dysphonia, is a voice disorder of neurological aetiology that affects the laryngeal muscles causing intermittent spasms.^1^ These spasms prevent the vocal folds from vibrating efficiently and result in involuntary interruptions, but only during voiced speech and not during other vocalisations like coughing and laughing,^2^ making it a focal or task-specific dystonia.^1^ LD affects approximately one in 100,000 people^3–5^ and is a chronic condition with largely unknown causes. More than 80% of patients with LD have the adductor type of LD, in which the muscle spasms bring the vocal folds together, causing them to shut too tightly and thus disrupting the initiation of voicing.^1, 6–8^ A less common form of LD is the abductor type, in which the muscle spasms push the vocal folds apart from each other, resulting in overly breathy voice. Currently, treatment options for LD are limited^9^ with most patients opting for temporary symptom relief through speech therapy and/or Botulinum toxin (botox) injections.^10–13^ However, since botox injections provide only temporary relief, patients have to schedule periodic clinical visits. Surgical nerve resection has been explored as a potential treatment^14^ but most patients experience a recurrence of symptoms due to nerve regrowth.^15^ Better understanding of the pathophysiology of LD may help us to improve existing treatments and may contribute towards the design of novel and perhaps more effective treatments for LD.

The task specificity of LD suggests it has a central nervous system (CNS) origin, and indeed studies have found that many regions of the CNS exhibit abnormalities in LD. Some studies have found structural abnormalities via post-mortem analysis^16^ or MRI-based morphometry measurements of grey matter volume, cortical thickness, cortical surface area and local white matter integrity.^17–21^ Others have found abnormal functional activity during speaking using PET^22^ or functional MRI (fMRI).^18, 23–25^ Taken together, these studies have found LD-associated abnormalities in: (1) areas that exhibit abnormalities across all dystonias: primary motor cortex (M1), primary somatosensory cortex (S1), thalamus, cerebellum and basal ganglia; (2) areas of the parietal and premotor cortices associated with task-specific dystonias^17^; (3) areas more specifically associated with speech motor control and speech processing^26^: e.g., inferior frontal gyrus (IFG), middle frontal gyrus (MFG), superior temporal gyrus (STG), middle temporal gyrus (MTG), and frontal operculum. The laryngeal motor cortex (LMC) in particular has been studied with a number of differing modalities. Studies of evoked response potentials^27^ and studies using transcranial magnetic stimulation^28, 29^ have generally shown hyper-excitability of LMC in LD. Notwithstanding the commonalities, the studies investigating abnormal activity during speaking in adductor LD, however, have shown conflicting findings (see Supp. Table 1 for a meta-analytical synopsis) with both increases and decreases in activity and connectivity within the speech motor control network in LD. For instance, during overt speech production, S1 shows hyperactivity in some studies^24, 25, 30^ and hypoactivity in others.^22, 23^ One possible source of these discrepancies is that many functional imaging studies use fMRI and PET which have poor temporal resolution.^31^ Vocal motor control is a dynamic process that involves the orchestration of various parts of the brain, respiratory muscles and speech articulators.^32^ There are a number of events related to onset of phonation and according to models of speech motor control,^33–36^ the different phonation events divide the phonation process into different time intervals which allow for functional interpretation of neural activity in those time intervals. Neural activity before initial glottal closure is associated with preparatory or related to feedforward control of speech.^33, 35^ Neural activity immediately after glottal closure, and prior to voice onset, also includes responses to somatosensory feedback. After the onset of phonation, neural activity also includes responses to the onset of auditory feedback. Thus, in addition to knowing where in the CNS aberrant activity occurs, it is also important to know during which vocal events (e.g. initial glottal closure, beginning of phonation or voice onset) abnormalities related to LD emerge because the sequence of events separates the phonation process into functionally interpretable time periods. At a more fundamental level, the insufficient temporal resolution provided by neuroimaging modalities like fMRI and PET cannot distinguish between activity related to the long-term *trait* of LD from the changes in neural activity related to moment-by-moment changes in the dystonic *state* of the larynx.

A neuroimaging modality with the temporal resolution needed to examine the rapid sequence of activations associated with phonation onset is magnetoencephalography (MEG). MEG in combination with advanced source reconstruction algorithms makes it possible to preserve spatial resolution while examining cortical activity on the order of milliseconds. In this study, we examined neural activity using MEG during the onset and continuation of phonation in patients with Adductor Laryngeal Dystonia (henceforth referred to as patients with LD) and compared it with neural activity in a control group. To the best of our knowledge, no prior study has looked at the time course of cortical activity prior to and during sustained phonation in LD (but see Khosravani *et al*.,^27^ for an analysis of spectral power using EEG during early vocalisation and late vocalisation). Since our intention was to investigate the temporal dynamics of neural activity in patients with LD, the high-gamma band (65-150Hz) with its high temporal resolution was a principal focus of our study. High-gamma band activity within sensorimotor cortices has been shown to be computationally related to speech motor planning and execution.^37, 38^ The larger windows of analysis needed for analysing lower frequency bands makes them less suitable for following rapid changes in neural activity. However, signals in beta band (12-30Hz) have been shown to play a role in motor planning, top-down motor-auditory interactions and the production of overt speech.^39–41^ Thus, we also examined beta-band activity in this study.

Furthermore, to isolate whether LD involves deficits in auditory feedback processing, we also briefly perturbed the pitch of participants’ auditory feedback during sustained phonation. Studies have shown that speakers respond to this auditory feedback perturbation by making compensatory changes to the pitch of their speech output.^42, 43^ Thus, perturbing pitch feedback not only allows us to test vocal behavioural responses to an altered auditory feedback but also examine neural activity induced by the onset of perturbation. Our previous studies have shown that pitch perturbations affect high-gamma-band oscillations.^44, 45^

In sum, the study design enabled us to make the following functional inferences about abnormal activity observed in the CNS in LD during phonation:

1. If observed in the premotor and motor cortices prior to glottal movement onset, it would suggest impairment in the processes involved in preparing to phonate.
2. If observed in somatosensory cortex after glottal movement onset, it would suggest impairments in somatosensory feedback processing.
3. If observed in auditory cortices after voice onset, it would suggest impairments in auditory feedback processing.
4. If observed in premotor, motor, somatosensory and auditory cortices after pitch perturbation onset, it would suggest impairments of these brain regions’ involvement in the auditory feedback control of vocal pitch.

## Materials and methods

### Participants

In this study, 22 patients (15 female, mean age = 57.38 years, standard deviation = 9.69 years) were recruited from the University of California, San Francisco (UCSF) Voice and Swallowing Center and through postings on the National Spasmodic Dysphonia Association’s website. Additionally, 18 controls (eight female, mean age = 53.25 years, standard deviation = 16.22 years) were recruited by word-of-mouth and from healthy research cohorts at the UCSF Memory and Aging Center. Across the groups with this sample size, for a power of 0.9, a generalized unpaired t-test with similar variance will detect effect sizes of 1.05. A two-sample heteroscedastic t-test was used to determine that the cohorts did not differ in age (*p* = 0.351, Table 1). The imaging part of the study (MEG and structural MRI acquisition) included 17 patients and 13 controls who also underwent a thorough clinical voice evaluation. Five patients participated only in the pitch perturbation task without imaging. High-gamma band data from five controls who participated in the pitch perturbation task along with MEG, was included to improve the signal-to-noise ratio specifically for high-gamma-band analysis locked to pitch perturbation onset. MEG data from two patients and two controls had to be excluded because of large dental or movement artefacts. Vocal behavioural data for five of the 22 patients had to be excluded because of poor pitch tracking. All patients were diagnosed by a team of laryngologists. Two of the 22 patients were also diagnosed to have vocal tremor. For a description of how many patients and controls were included in each analysis, see Supp. Table 2. Eligibility criteria for patients were: 1. a diagnosis of adductor LD, 2. symptomaticity during research participation, 3. at least three months since their previous botox injection. Eligibility criteria for control participants were: 1. no structural MRI abnormalities, 2. no hearing loss and 3. absence of neurological disorders. The study was approved by the Committee on Human Research of UCSF. Informed consent was obtained from each participant prior to the study.

### MRI Acquisition

T1-weighted structural MRI images were acquired using a 3-Tesla MRI scanner (Discovery MR750, GE Medical Systems) with an 8-channel head coil at the UCSF Margaret Hart Surbeck Laboratory for Advanced Imaging. An inversion recovery spoiled gradient echo sequence was used to acquire 128 axial slices (repetition time = 6.664ms, echo time = 2.472ms, inversion time = 900ms, slice thickness = 1mm, in-plane voxel dimensions = 0.5mm x 0.5mm). Individual structural MRIs were spatially normalised to a standard Montreal Neurological Institute (MNI) template using SPM8 (https://www.fil.ion.ucl.ac.uk/spm/software/spm8/).

### MEG Imaging and Experimental Design

The MEG scanner (CTF, Coquitlam, BC, Canada) consisted of 275 axial gradiometers. Participants were scanned in supine position and signals were collected at a sampling rate of 1200Hz. Head position was recorded relative to the sensor array using three fiducial coils placed at the nasion and left and right preauricular points. These points were co-registered with individual structural MRI images to generate head shapes.

The experimental task consisted of 120 trials. During each trial, participants were prompted to start vocalising the vowel /ɑ/ upon seeing a green dot on a screen (Fig. 1). They were asked to hold the phonation for the duration of the visual prompt (∼2.4s). Pre-phonatory laryngeal muscular activity (glottal movement onset) was recorded using surface EMG electrodes which detect intrinsic laryngeal muscular activity as reliably as needle EMG electrodes.^46^ An abrasive gel was used to prepare the skin and to lower the impedance. Two electrodes were then pasted on either side of the larynx over the cricoid ring and a third electrode was placed over the thyrohyoid membrane. EMG signals were recorded using the double differential technique.^47^ Conductive paste was used to increase conductivity between the skin and the electrodes. The ground electrode was placed on the participant’s forehead. Simultaneous electrocardiography (ECG) and electrooculography (EOG) signals were also collected to ensure that the electrophysiological signals picked up by the EMG electrodes were devoid of noise from eye blinks, eye movements and heartbeats. Participants’ vocal output was picked up by an optical microphone (Phone-Or Ltd., Or-Yehuda, Israel), passed through a digital signal processing system (DSP) and fed back to them via insert earphones (ER-3A, Etymotic Research, Inc., Elk Grove Village, IL). On every trial, between 200ms and 500ms after voice onset, the DSP perturbed the pitch of their auditory feedback by either +100 cents or -100 cents (1/12^th^ of an octave) for 400ms and vocal responses to this change were recorded. The DSP was implemented on a PC as a real-time vocoder program called Feedback Utility for Speech Processing (FUSP).^43, 45, 48, 49^

**Figure 1.**
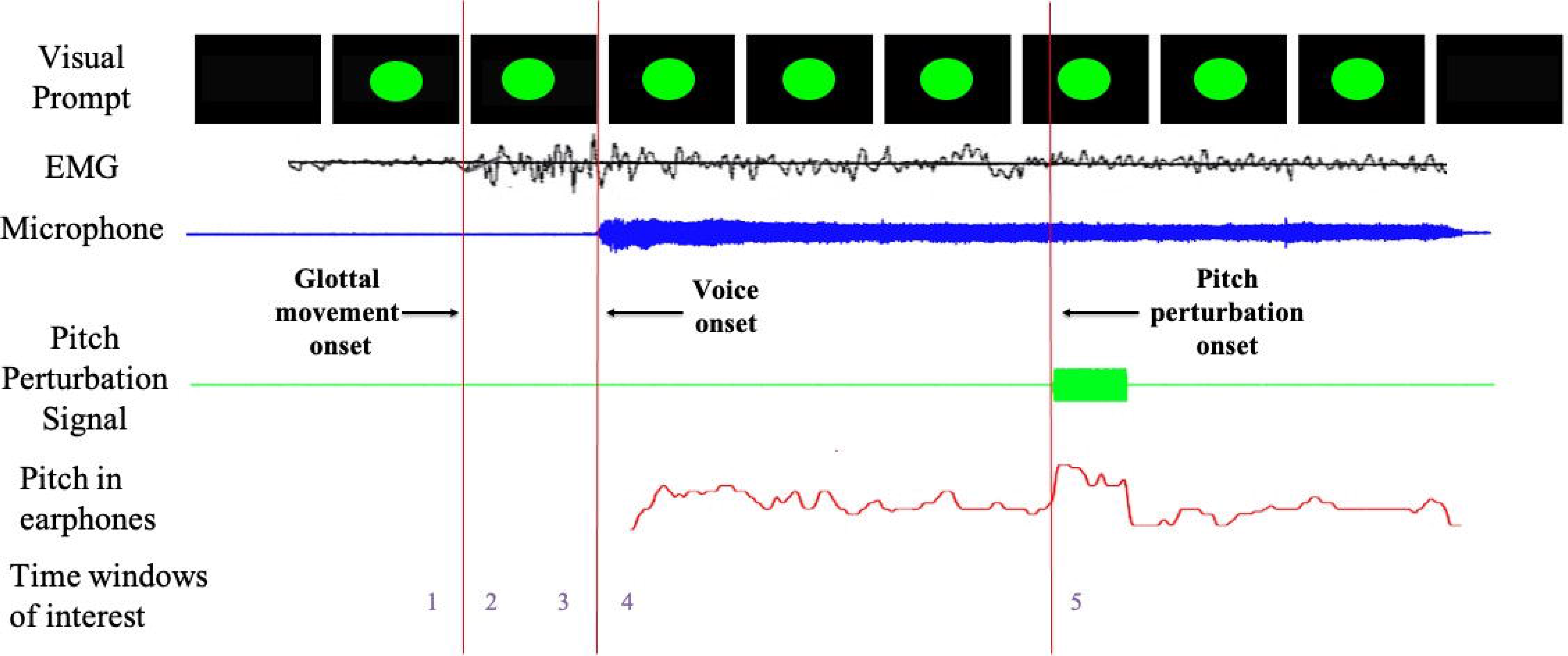
Schematic diagram of the task. Participants were prompted to vocalise the vowel /ɑ/ into a microphone for as long as they saw the green dot on the display. Participants could hear themselves through earphones throughout every trial. Glottal movement onset was measured using surface electromyography. Subjects’ pitch was altered using a digital signal processing unit, between 200-500ms after voice onset, either up or down by 100 cents (1/12^th^ of an octave) for 400ms and sent this shifted signal to the participants’ earphones. The numbers at the bottom of the figure represent various time windows as follows: 1 = before glottal movement onset, 2 = after glottal movement onset, 3 = before voice onset, 4 = after voice onset, 5 = response to pitch perturbation onset.

### Data Analysis

#### Pitch Data Processing

Audio signals for participants’ speech and the altered feedback were both collected at 11,025 Hz. The pitch time course for phonation in each trial was determined using an autocorrelation-based pitch tracking method.^50^ Pitch tracks for all the trials were extracted and were aligned from 200ms before perturbation onset to 1000ms after perturbation onset. Trials with pitch tracking errors or with incomplete utterances were marked bad and excluded. For the remaining good trials, pitch was converted from Hertz to cents using the following formula:

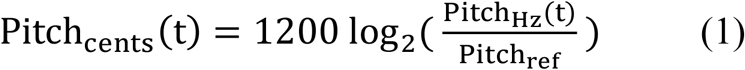

where Pitch_Hz_(t) is the pitch in Hertz at timepoint t and Pitch_ref_ is the reference pitch calculated as the mean pitch during a window spanning from 50ms prior to perturbation onset to 50ms after perturbation onset. Participants responded to the feedback perturbation by deviating from their baseline pitch track. For each participant, responses to both upward and downward perturbations were calculated and pooled together. To do this, the responses to upward perturbations were flipped and combined with responses to downward perturbations. In this scheme, all compensatory responses are positive and all following responses are negative, irrespective of the direction of the applied shift. Mean responses for patients and controls were plotted.

Additionally, peak deviation from the baseline pitch track was calculated for every trial. This peak deviation had a positive sign if the response was opposing the shift (or compensatory) and negative if the response was following the shift. The mean across trials of this peak deviation was called the mean compensation to pitch perturbation for each participant.

#### Correlations between responses to pitch perturbation and voice evaluation measures

To examine whether compensatory responses to pitch perturbation can predict disease severity, we selected three measures of voice impairment (Table 1) from the participants’ clinical voice evaluation: i) A subjective measure: Voice Handicap Index (VHI), a validated 30-item self-reported voice assessment,^51^ ii) A perceptual measure: Consensus Auditory-Perceptual Evaluation of Voice (CAPE-V)^52^ and a iii) A laryngeal motor control measure: the Laryngeal Diadochokinesis rate (L-DDK rate).^53–55^ To calculate the L-DDK rate, participants were asked to repeat the vowel / ʌ / as fast and controlled as they could, while their voice was being recorded, until they were asked to stop by the Speech and Language Pathologist. Their speech signal was then edited to display 7 seconds of utterances and the L-DDK rate was calculated as the number of syllables uttered per second.

#### MEG Data Processing

Third-order gradient noise correction and Direct Current offset correction were performed on the MEG datasets. A notch filter was implemented at 120Hz (width = 2Hz) to reduce power line noise. EMG signals, voice signals and MEG signals were examined simultaneously, and three markers were added to mark the onset of glottal movement, onset of voicing and onset of pitch perturbation. Subsequent neural analyses were focused on estimating event-related non-phase-locked activity with respect to these phonatory event markers. Trials with abnormal signals due to head movement, eye blinks or saccades were excluded. Spatiotemporal source localisation for induced neural activity was reconstructed for each individual subject and then mapped on to the spatially-normalised structural MRI for that subject using time-frequency-optimised adaptive spatial filtering (8mm lead field) in the Neurodynamic Utility Toolbox for MEG (NUTMEG: http://nutmeg.berkeley.edu).^56, 57^ This source space reconstruction provided a voxel-by-voxel estimate of neural activity derived using a linear combination of a spatial weight matrix and a sensor data matrix.

#### Phonatory onset interval analysis

Using the glottal movement onset marker and the voice onset marker added during the MEG analysis, phonatory onset interval was calculated for every trial in every participant to look at differences in the mean phonatory onset interval in patients as compared to controls, which has previously been shown to be elongated in patients with LD.^58^

### Time windows of interest

To better interpret the differences in activity between patients and controls, the entire time course of the trial was divided into six time windows around the three phonatory event markers (Fig. 1).

• Time window 1: 125ms preceding glottal movement onset.
• Time window 2: 125ms succeeding glottal movement onset.
• Time window 3: 125ms preceding voice onset
• Time window 4: 125ms succeeding voice onset
• Time window 5: responses to pitch perturbation onset.

### Statistical analysis

For time windows showing qualitative differences in behavioural responses to pitch perturbation, a linear mixed effects model was run in SAS 9.4 (SAS Institute Inc., Cary, NC) with group as the independent variable and pitch response in cents as the dependent variable.

To account for multiple timepoints from each participant’s mean vocal pitch response time-course, participant identity was included as a repeated measure with a random intercept.

To look for correlations between responses to pitch perturbation and voice evaluation measures, three Pearson’s correlation coefficients were calculated between participants’ mean compensation to pitch perturbation and VHI, CAPE-V and L-DDK rate.

For neural data analysis, active windows were defined as the time period after each of the three phonatory event markers. Noise-corrected pseudo-F statistics were computed by comparing the active window to a control window (pre-stimulus for glottal movement onset and voice onset analysis, pre-perturbation for perturbation onset analysis). Within and between-group statistical analyses were performed using statistical non-parametric mapping methods incorporated into the NUTMEG toolbox.^59^ For voice onset analysis, the phonatory onset interval or the duration between glottal movement onset and voice onset was included as a covariate in the statistical model. To correct for multiple comparisons across time and space, corrected p-value thresholds were calculated for *α* = 0.05 and a False Discovery Rate (FDR) of 5%. Furthermore, cluster correction was performed to exclude clusters with less than 18 congruent voxels as done in previous studies.^48^

To look at differences in mean phonatory onset interval between patients and controls, a two-sample heteroscedastic t-test was run.

### Data availability

The data that supports the findings of this study will be made available upon reasonable request.

## Results

### Initiation of voice production

#### Larger phonatory onset interval in patients with LD

Analysis of the time interval between glottal movement onset and voice onset markers (time windows 2 and 3) in the current study showed (Fig. 2) a significantly larger duration (*p* = 1.14 x 10^-47^; two-sample heteroscedastic t-test) in LD patients (*n* = 15, *mean* = 356.5ms, *standard error of the mean* = 4.94ms) than in controls (*n* = 11, *mean* = 251.55ms, *standard error of the mean* = 4.1ms). These results serve as a replication of a previous study showing voice onset delay in patients with LD.^58^

**Figure 2.**
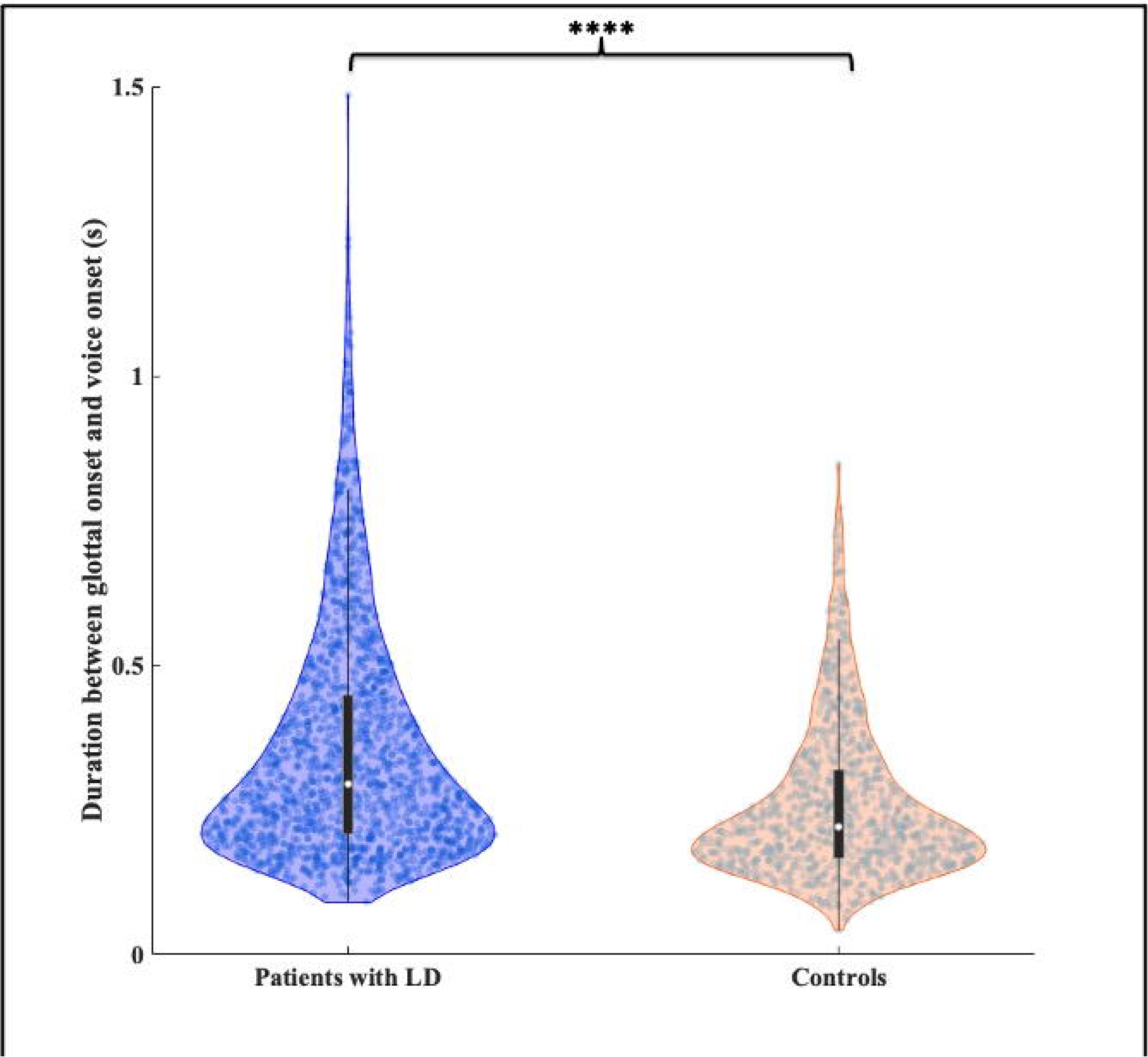
Phonatory onset interval. Patients with LD have a larger phonatory onset interval (duration between glottal movement onset and voice onset) as compared to controls (p = 1.140 x 10^-47^; two-sample heteroscedastic t-test). Data points within the violin plots represent phonatory onset interval time for individual trials. Boxplots within the violins indicate the median (white dot) and interquartile range (IQR), the whiskers indicate 1.5 * IQR.

#### Bilateral reduction in inferior frontal and increase in parietal beta-band activity around glottal movement onset

Widespread differences in beta-band neural activity between controls and patients were observed before and after glottal movement onset (Fig. 3A, Supp. Table 3). Even prior to glottal movement (time window 1), the greatest reductions in beta-band activity in patients were seen in the left ventral motor cortex (vMC), left ventral premotor cortex (vPMC) and left inferior frontal gyrus (IFG). The reduced activity in left vPMC and left IFG persisted after glottal movement onset (time window 2), but differences were smaller after onset. Reduced activity was also observed in the anterior left STG, anterior left MTG, right middle frontal gyrus (MFG), right cerebellum and right IFG after glottal movement onset. In contrast, patients with LD also showed persistently increased activity, both prior to and after glottal movement onset, bilaterally in a large cluster in the inferior parietal lobule (IPL). Increased activity was also observed in right dorsal premotor cortex from -25ms before glottal movement onset to +125ms after and left dorsal premotor cortex at +125ms after glottal movement onset.

**Figure 3.**
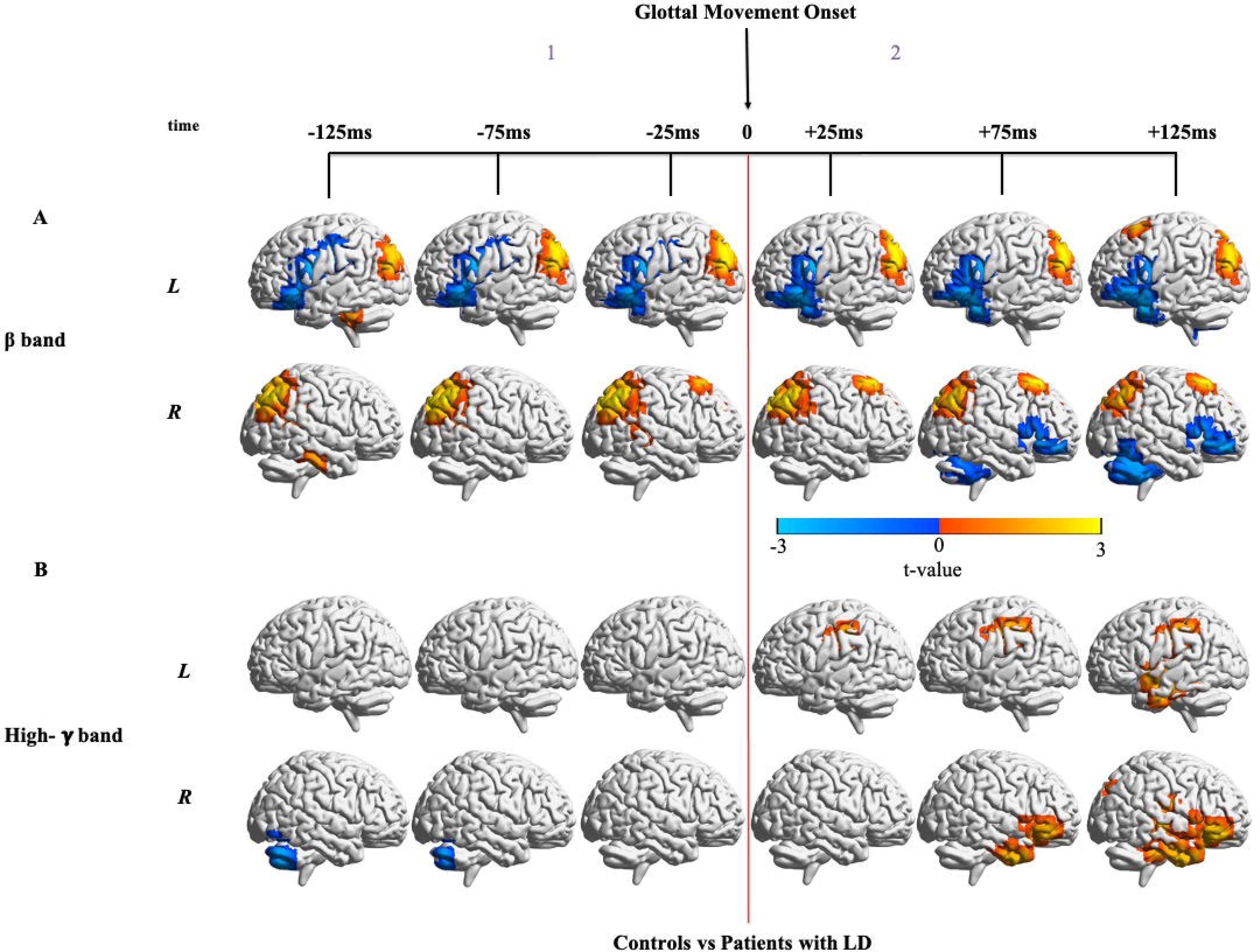
Differences in neural activity around glottal movement onset between controls and patients with LD. (**A)** Non-phase-locked beta-band activity (12-30Hz) differences between patients and controls locked to glottal movement onset. As compared to controls (n = 11), patients with LD (n = 15) show significant differences in beta-band activity both before and after glottal movement onset. False Discovery Rate (FDR) correction for a rate of 5% and cluster correction at a threshold of 18 voxels and p < 0.05 were performed. For beta-band neural activity locked to glottal movement onset in each group alone, refer to Supplementary Fig. 1. (**B)** Non-phase-locked high-gamma-band activity (65-150Hz) differences between patients and controls locked to glottal movement onset. As compared to controls (n = 11), patients with LD (n = 15) show significantly increased activity after glottal movement onset. False Discovery Rate (FDR) correction for a rate of 5% and cluster correction at a threshold of 18 voxels and p < 0.05 were performed. For high-gamma-band neural activity locked to glottal movement onset in each group alone, refer to Supplementary Fig. 4.

#### Reduced right cerebellar activation before and increased bilateral cortical activation after glottal movement onset in high-gamma band

Differences in high-gamma-band neural activity between controls and patients showed an interesting change in polarity from before glottal movement onset to after (Fig. 3B, Supp.Table 4). From -125 to -75ms before glottal movement onset (time window 1), patients showed reduced activity in the right cerebellum. However, after glottal movement onset (time window 2), patients showed increased activity in the left primary somatosensory cortex (postcentral gyrus/S1) from +25 to +125ms. Increased activity in patients was also observed in the right anterior temporal lobe (ATL) and right IFG from +75 to +125ms after glottal movement onset. At +125ms after glottal movement onset, patients also showed increased high-gamma-band activity bilaterally in the STG and MTG.

#### Bilateral increase in dorsal sensorimotor cortical activation and reduced cerebellar activation in beta band around voice onset

Prior to voice onset in beta band (time window 3), bilateral hyperactivity in patients’ IPLs and hypoactivity in their left IFG and anterior left STG and MTG was observed (Fig. 4A, Supp. Table 5), reflecting differences persisting after glottal movement onset (time window 2) observed in Fig. 3A. Importantly, both before and after voice onset, patients showed increasing bilateral hyperactivity along S1 and the superior parietal lobule (SPL). Patients also showed hyperactivity in the superior left MFG from -125ms to +25ms. In contrast, patients showed reduced activity bilaterally in the cerebellum and inferior occipital lobe (greater differences in the right hemisphere) after voice onset (time window 4). Patients also showed reduced activity in the right MFG and right IFG after voice onset. Note that these results were not impacted by differences in phonatory onset interval between the two groups, because this interval was included as a covariate in the statistical analysis.

**Figure 4.**
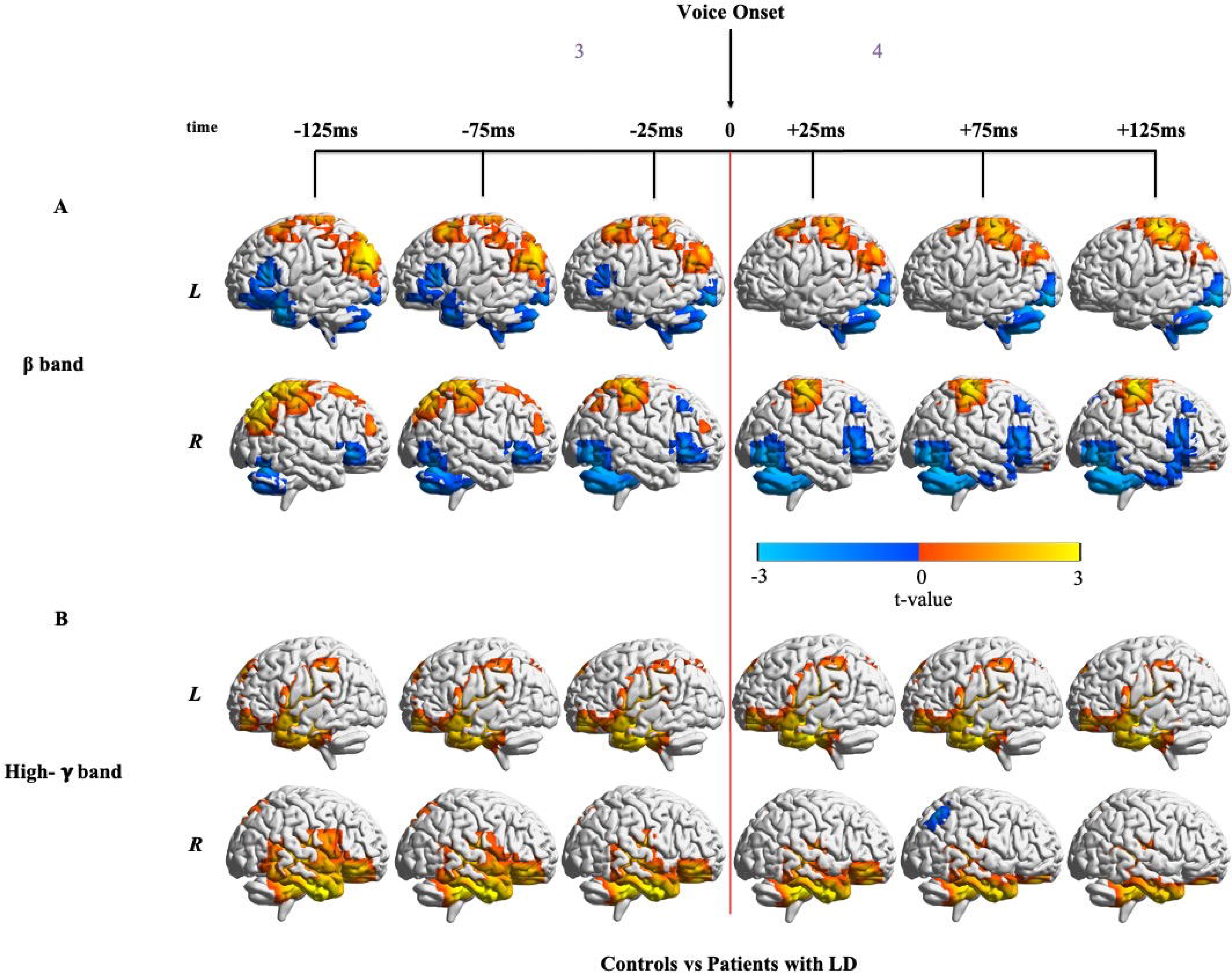
Differences in neural activity around voice onset between controls and patients with LD (A) Non-phase-locked beta-band activity (12-30Hz) differences between patients and controls locked to voice onset. Phonatory onset interval was added as a covariate in the statistical analysis. As compared to controls (n = 11), patients with SD (n = 15) show significant differences both before and after voice onset. False Discovery Rate (FDR) correction for a rate of 5% and cluster correction at a threshold of 18 voxels and p < 0.05 were performed. For beta-band neural activity locked to voice onset in each group alone, refer to Supplementary Fig. 2. (B) Non-phase-locked high-gamma-band activity (65-150Hz) differences between patients and controls locked to voice onset. Phonatory onset interval was added as a covariate in the statistical analysis. As compared to controls (n = 11), patients with SD (n = 15) show consistent differences in both hemispheres from before voice onset through voice onset. These differences increase after voice onset in the left hemisphere and decrease in the right hemisphere. False Discovery Rate (FDR) correction for a rate of 5% and cluster correction at a threshold of 18 voxels and p < 0.05 were performed. For high-gamma-band neural activity locked to voice onset in each group alone, refer to Supplementary Fig. 5.

#### Bilateral increase in activation in ventral sensorimotor cortex, prefrontal cortex and temporal lobe in high-gamma band around voice onset

Patients showed widespread hyperactivity in high-gamma band both before and after voice onset (Fig. 4B, Supp. Table 6) bilaterally in the ventral sensorimotor cortex, temporal lobe, medial and ventral parts of the left prefrontal cortex and right ventral prefrontal cortex. Hyperactivity in the left temporal lobe increased from before voice onset (time window 3) to after voice onset (time window 4) whereas hyperactivity in the right temporal lobe decreased with time. Hyperactivity was also observed in patients in the right precuneus from -125 to -75ms before voice onset (time window 3). Patients showed hypoactivity in the right IPL at

+75ms after voice onset (time window 4).

### Response to pitch feedback perturbation

#### Pitch perturbation vocal responses do not differ in patients with LD

No statistical differences were found between patients’ and controls’ vocal responses to pitch perturbation (Fig. 5A). Although there appeared to be a slight reduction in patients’ vocal response from 250 to 400ms after pitch perturbation onset, this reduction was not significantly different from controls’ vocal response in the same window [LMM, *F*(1, 2001) = 0.01, *p* = 0.909]. To better understand the neural mechanism driving the response to pitch perturbation onset, we decided to further split time window 5 into two parts, 5early (from 0 to 225ms after pitch perturbation onset) and 5late (from 225 to 450ms after pitch perturbation onset).

**Figure 5.**
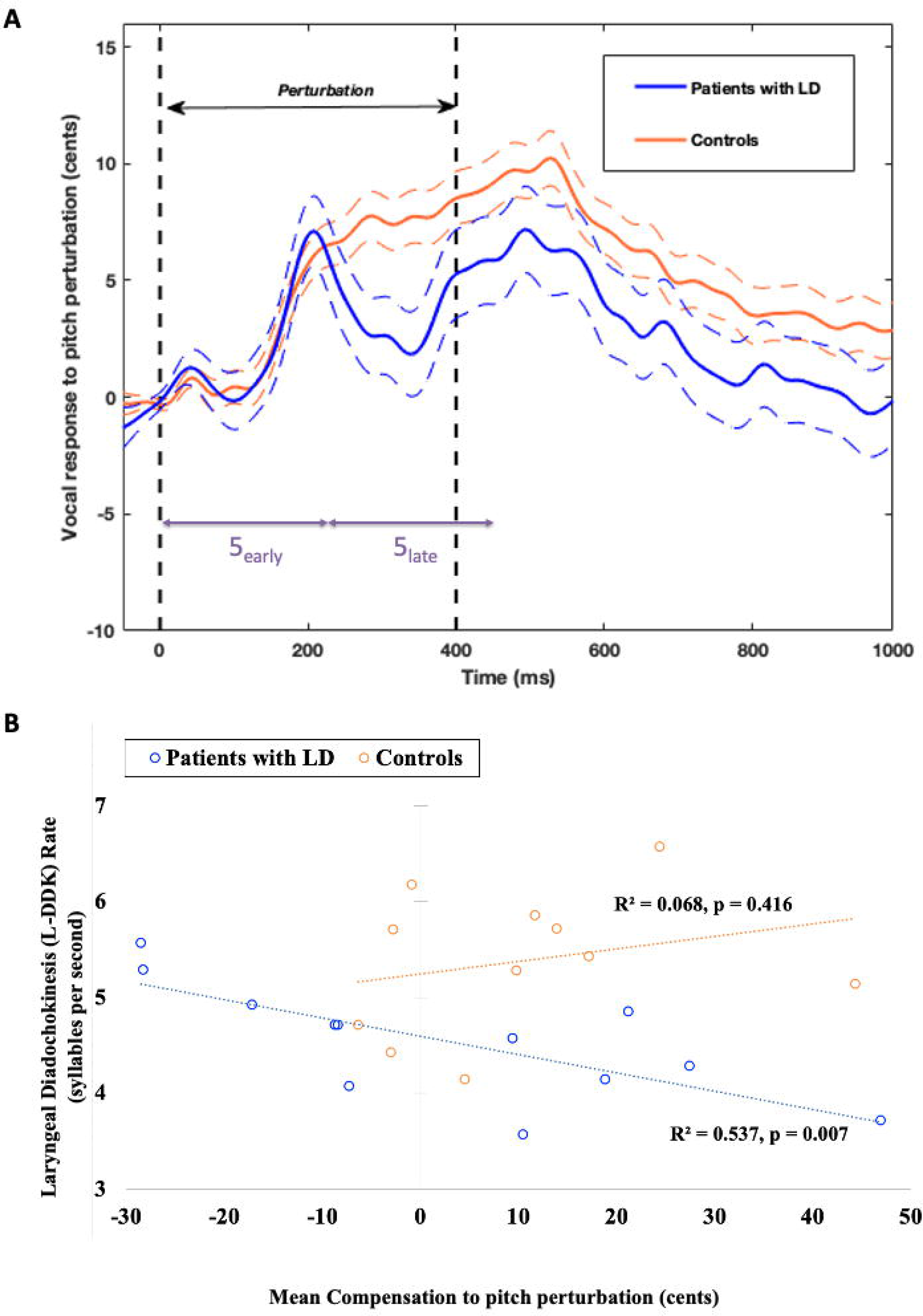
Pitch control in patients with LD and controls (A) Vocal response to pitch perturbation: Patients with LD (n = 17) have a response to pitch perturbation that is not statistically different from that in controls (n = 12) [LMM, *F*(1, 2001) = 0.01, *p* = 0.909]. The solid lines are the mean responses and the flanking dashed lines indicate standard error of the mean responses. Time windows 5a and 5b represent early and late response to pitch perturbation **(B)** Correlation between mean compensation and laryngeal diadochokinesis (L-DDK) rate: In patients with LD, the L-DDK rate is negatively correlated with mean compensation to pitch perturbation. Patients (n = 12) who have a higher syllable rate tend to follow the direction of the pitch shift and the ones with a lower syllable rate tend to have a larger compensation value for pitch shifts. This correlation does not hold true in controls (n = 11).

Patients showed larger pitch variability in cents both within-trial (*p* = 0.008; two-sample heteroscedastic t-test) and across trials (*p* = 0.02; two-sample heteroscedastic t-test) in a 200ms pre-perturbation time window, i.e., time window 4 (Supplementary Fig. 1) indicating that patients not only possessed the vocal range to compensate for pitch perturbation but also had greater variations in laryngeal control during sustained phonation, perhaps owing to the presence of spasms.

#### Compensation to pitch perturbation in LD predicted severity of disease

In patients, mean compensation was negatively correlated (Fig. 5B) with L-DDK rate (*R^2^* = 0.537, *p* = 0.007), but no correlations were found, in both patients and controls, between mean compensation and VHI (patients: *R^2^* = 0.004, *p* = 0.839; controls: *R^2^* = 0.150, *p* = 0.239) or mean compensation and CAPE-V Overall Severity (patients: *R^2^* = 0.037, *p* = 0.549; controls: *R^2^* = 0.016, *p* = 0.709). Patients with lower L-DDK rates, i.e., with greater disease severity, had greater values of mean compensation. Patients with higher L-DDK rates, i.e., with lesser disease severity, either had smaller values of mean compensation or tended to follow the direction of the pitch shift. Notably, in controls, there was no correlation between mean compensation values and L-DDK rates (*R^2^* = 0.068, *p* = 0.416).

#### Bilateral increased frontoparietal beta-band activity after pitch perturbation onset

Examination of beta-band neural activity after pitch perturbation onset (Fig. 6A, Supp. Table 7) showed that patients had greater activity bilaterally in the IFG and in right vPMC and vMC with activity peaking from 125 to 225ms after perturbation onset. Frontal hyperactivity in the right hemisphere was more widespread than in the left hemisphere. Patients also showed greater activity in the left SPL from 175 to 375ms, right IPL from 375 to 425ms, right superior frontal gyrus (SFG) from 225 to 425ms and anterior right STG from 25 to 425ms. Although the parietal hyperactivity was bilateral, the left hemisphere led the right hemisphere in terms of time of activity increase. Cerebellar activations were mixed – there was early left hemisphere reduction (time window 5_early_) and right hemispheric activity increased (time window 5_late_) paralleling the increased activity in the right frontal lobe. Note that neural differences in beta band were observed despite no behavioural differences in vocal pitch responses to pitch feedback perturbations.

**Figure 6.**
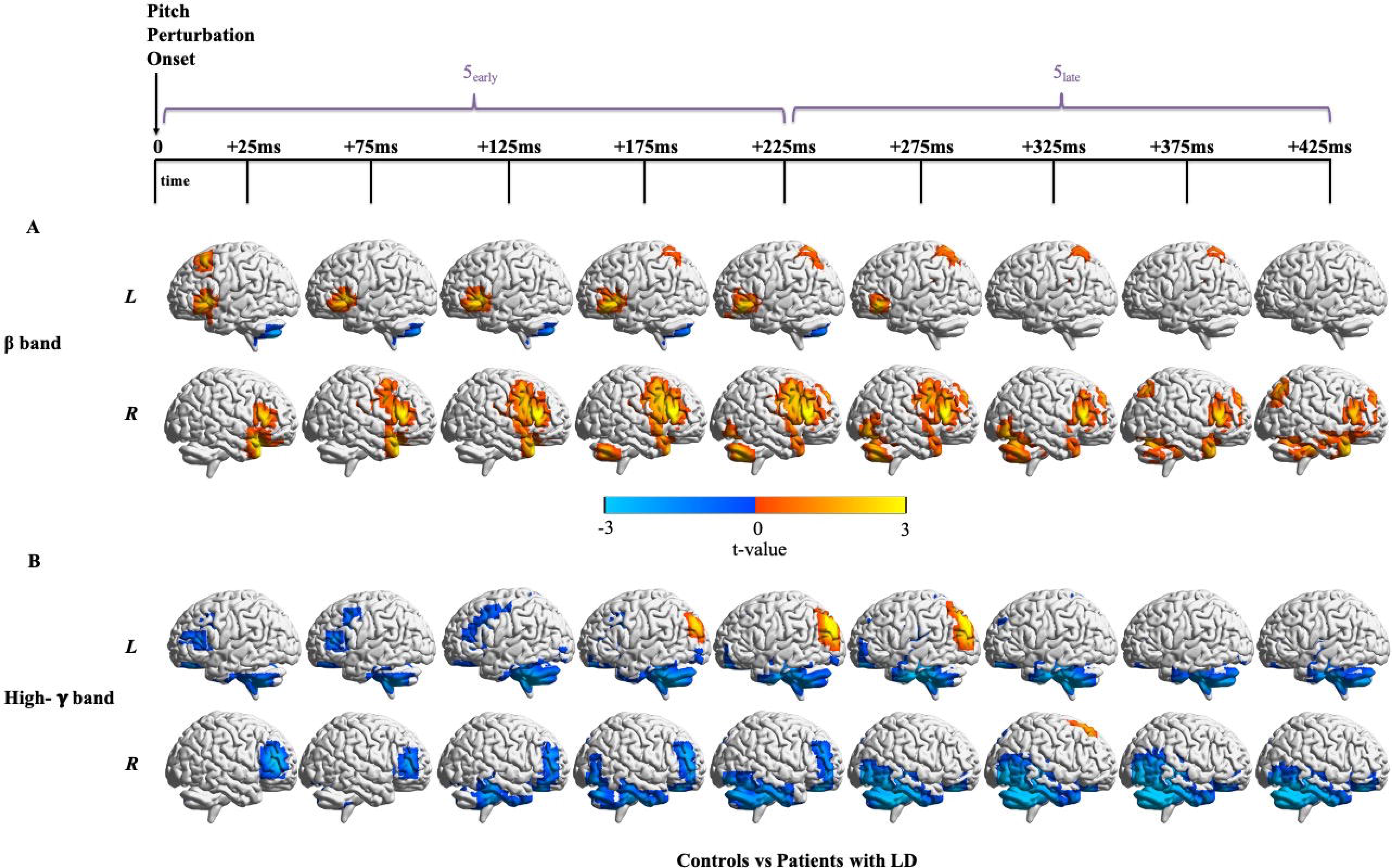
Differences in neural activity around pitch perturbation onset between controls and patients with LD (A) Non-phase-locked beta-band activity (12-30Hz) differences between patients and controls locked to pitch perturbation onset. Patients with LD (n = 15) show greater beta-band activity as compared to controls (n = 11) in a number of regions. False Discovery Rate (FDR) correction for a rate of 5% and cluster correction at a threshold of 18 voxels and p < 0.05 were performed. For beta-band neural activity locked to pitch perturbation onset in each group alone, refer to Supplementary Fig. 3. (**B)** Non-phase-locked high-gamma-band activity (65-150Hz) differences between patients and controls locked to pitch perturbation onset. Patients with LD (n = 15) show mostly lesser high-gamma-band activity as compared to controls (n = 16) along with greater activity in some regions. False Discovery Rate (FDR) correction for a rate of 5% and cluster correction at a threshold of 18 voxels and p < 0.05 were performed. For high-gamma-band neural activity locked to pitch perturbation onset in each group alone, refer to Supplementary Fig. 6.

#### Bilateral reduced cerebellar, prefrontal and temporal high-gamma-band activity after pitch perturbation onset

Patients showed widespread reduced high-gamma band activity after pitch perturbation onset (Fig. 6B, Supp. Table 8). Hypoactivity in the left cerebellum preceded that in the right cerebellum and peaked earlier. Bilateral hypoactivity in the MFG and IFG dissipated as time progressed whereas that in the ventral prefrontal cortex persisted from 25 to 425ms. There was significant hypoactivity in the left inferior temporal lobe from 175 to 375ms and from 125 to 425ms in the right temporal and occipital lobes. Hyperactivity in patients was also observed in the left IPL from 175 to 275ms after pitch perturbation onset and right dorsal SFG at 325ms. Again, note that neural differences in high-gamma band were observed despite no behavioural differences in vocal pitch responses to pitch feedback perturbations.

## Discussion

The current study used MEG imaging to determine whether patients with LD exhibited abnormal neural activity during the initiation of phonation and in response to pitch feedback perturbation. Non-phase-locked event-related beta and high-gamma band abnormalities were observed at different times in multiple regions within the speech motor control network consistent with electrophysiological recordings reflecting local neural population activity.^60–62^ Prior to glottal movement onset, patients showed reduced beta-band activity in the left inferior frontal cortex, increased beta-band activity in the parietal cortex and reduced high-gamma-band activity in the right cerebellum. After glottal movement onset, the reduced inferior frontal and increased parietal beta-band activity continued and patients also showed increased bilateral cortical activation in high-gamma band. Around voice onset, patients showed bilaterally-increased dorsal sensorimotor cortical beta-band activity, reduced cerebellar beta-band activity and bilaterally-increased ventral sensorimotor, prefrontal and temporal lobe high-gamma-band activity. In response to pitch feedback perturbation, although patients’ compensatory vocal responses were similar to controls, they showed increased bilateral frontoparietal beta-band activity and reduced bilateral cerebellar, prefrontal and temporal lobe high-gamma-band activity. Patients’ mean compensation to pitch perturbation predicted disease severity. Below we discuss and interpret these results in detail, how they relate to prior neuroimaging studies in patients with LD, and additionally provide a functional interpretation of the findings in the context of our model of speech motor control.^34, 35^

### Abnormalities around glottal movement onset

Prior to glottal movement onset, patients showed reduced activity in the left vPMC and left VMC, the part of the motor cortex that is involved in laryngeal articulatory control.^32, 63^ Hypoactivation in alpha band in the left motor cortex has been observed in a previous EEG study.^27^ Patients also showed hypoactivity in the right cerebellum before glottal movement onset in the high-gamma band. The cerebellum is implicated in various aspects of speech motor control.^64–69^ Decreased activity in the right cerebellum has been observed in prior fMRI studies in patients with LD involving overt speech production, respiration and vocalisation.^24, 25^ This abnormal activity in the motor network prior to glottal movement onset suggests impairments in motor preparation.

Patients showed persistent hyperactivity in the high-gamma band after glottal movement onset in the left ventral somatosensory cortex and left supramarginal gyrus. Abnormal activation in S1 has been observed in previous studies^25^ and the current results highlight the temporal specificity of this abnormality. Abnormal activity in the sensorimotor cortices immediately following glottal movement onset suggests impaired sensitivity to somatosensory feedback processing.

Patients exhibited hyperactivity later in the right ATL, right IFG, bilateral STG and MTG. Abnormally increased activity in the STG and MTG has been reported previously in symptomatic and asymptomatic tasks in patients with LD.^25^ The activity in auditory processing regions prior to voice onset is surprising. One possibility is that the observed activity represents abnormalities in the generation of auditory expectations prior to voice onset.

### Abnormalities around voice onset

Beta-band hypoactivity was observed in left vPMC only prior to voice onset. In contrast, beta-band hyperactivity around the IPL was observed bilaterally prior to voice onset and persisted only in left IPL after voice onset. Bilateral high-gamma-band hyperactivity was observed along the temporal lobes prior to voice onset and continued even after voice onset. Both beta-band and high-gamma-band hyperactivity in patients in S1 that began prior to voice onset continued after voice onset. This somatosensory hyperactivity is a persistent signature in most previous studies.^24, 25, 27, 30^ The IPL has also been implicated in patients with LD in studies investigating resting-state functional network connectivity and functional activity during symptomatic speech production.^18, 70^ Since hyperactivity in somatosensory feedback processing is also seen at the poor temporal resolution of fMRI in both symptomatic and asymptomatic tasks,^24, 25, 30^ it may be a general trait of patients with LD. It is also notable that the beta-band hyperactivity in S1 is not seen in the primary auditory cortex (A1) and other higher order regions involved in the processing of auditory feedback (posterior superior temporal sulcus (pSTS) and ventral supramarginal gyrus (vSMG)). However, hyperactivity in the high-gamma band is observed in regions involved in auditory feedback processing both before and after voice onset. These abnormalities point to both somatosensory and auditory feedback processing deficits in LD.

Behaviourally, when we consider the transition between glottal closure and voice onset, we note that this transition took longer for patients. This elongation in the phonatory onset interval has also been observed in previous studies.^58^ One possible reason for the larger phonatory onset interval in patients with LD may be due to more time needed to build up subglottal pressure sufficient to overcome their tightly adducted vocal folds. Evidence for increased phonatory effort in these patients before and after voice onset is consistent with beta-band hyperactivity around the truncal sensorimotor cortex, which could perhaps indicate that the respiratory muscles work harder in patients with LD to push air out of a tightly closed glottis due to dystonia of the adductor muscles, thus perhaps reflecting momentary state characteristics of LD.

### Abnormalities around pitch perturbation onset

We isolate auditory feedback processing abnormalities by examining neural responses to auditory feedback perturbations. The current study is the first to examine neural responses to pitch perturbation in patients with LD. There was beta-band hyperactivity in patients, especially in the right vPMC and vMC regions known to be responsive to auditory feedback perturbations^71^. Patients also showed widespread hypoactivity in the high-gamma band after pitch perturbation onset, notably bilaterally in the cerebellum and the frontal and temporal lobes. Reduced activity has been observed in these regions in patients with LD during prolonged vowel phonation and reading tasks.^23, 24^ Post-perturbation-onset increased activity, despite a lack of behavioural differences in the compensatory vocal response, may be indicative of an increased effort in vocal control required to compensate for the auditory mismatch.

### Behavioural responses to pitch feedback perturbations

Behaviourally, in the current study, patients and controls showed similar compensatory vocal responses, in spite of significant neural activation differences. Patients’ mean response exhibited two peaks but this behaviour was not observed for each patient. In contrast to the current behavioural findings, a recent study found that patients with LD have a larger and earlier vocal pitch compensation response to pitch feedback perturbation.^72^ However, there are a few notable differences in the stimuli between the two studies; the pitch shift in Thomas *et al.*^72^ was 200 cents (100 cents here) and lasted for 200ms (400ms here). The patients participating in our study were all pre-botox whereas Thomas *et al.*^72^ recruited patients before and after botox. Nevertheless, discrepancies in behavioural findings between the current study and those of Thomas *et al.*^72^ warrant further follow-up in larger LD cohorts. We also found that patients had a significantly lower L-DDK rate and that their mean compensatory response to pitch feedback perturbation negatively correlated with their L-DDK rate. Taken together, the behavioural and neural findings in response to pitch feedback perturbations, and their associations with symptoms in LD suggest not only greater vocal effort required for pitch feedback control but also a different mode of pitch control in patients.

### Functional interpretations of neuroimaging abnormalities in LD

We explore functional interpretations of our results with reference to our SFC model of speech motor control. Many operations of SFC have their analogues in other models like DIVA.^33, 73, 74^ The SFC model of phonation (Fig. 7) is based on the idea that at any moment in time, the larynx has a laryngeal motor state and that depending on the state, it can produce either somatosensory feedback (like at glottal movement onset) or both somatosensory and auditory feedback (like at voice onset). When a speaker decides to phonate, the higher frontal cortex activates a phonation control network consisting of, but not limited to, the motor cortex, the somatosensory cortex and the auditory cortex. Activation of this network causes M1 to output laryngeal controls which initiate glottal movement (and thus the generation of somatosensory feedback) and an efference copy of these controls drives the vPMC, in conjunction with the cerebellum, to update its prediction of the current vocal tract state which is used to predict somatosensory feedback from the larynx. Any abnormalities in neural activity seen before this point would have to do with the output of motor cortex and the higher frontal cortex (IFG) and the initiation of feedback predictions by the premotor cortex. Once the somatosensory feedback reaches S1, it is compared with the predicted feedback. Any mismatch with the prediction results in corrections to the estimated laryngeal state in the vPMC, which in turn further modifies the output of controls by M1. The onset of phonation generates both somatosensory and auditory feedback. Like somatosensory feedback, auditory feedback is also compared with a feedback prediction. Any mismatch with the auditory prediction results in corrections to the estimated state and output of laryngeal controls. Thus, any abnormalities seen after voice onset would indicate abnormal responses to either the somatosensory or auditory feedback of phonation. In addition, experimental perturbation of the pitch of auditory feedback creates a deliberate mismatch between the prediction and the actual auditory feedback and generates auditory feedback prediction errors in the auditory cortex. This error usually causes a compensatory vocal response driven by M1 that shifts vocal pitch production in the direction opposite to the perturbation.

**Figure 7.**
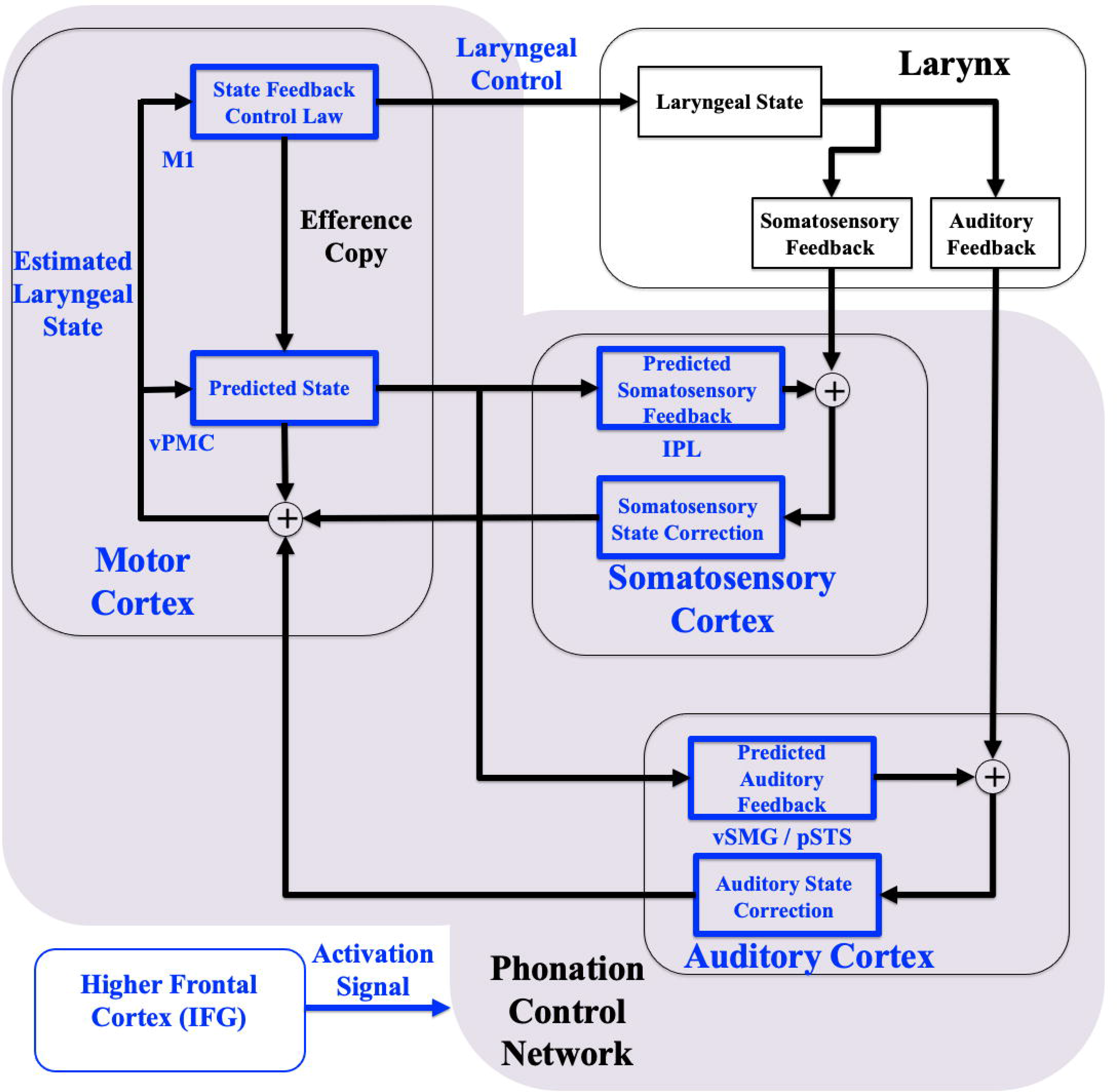
Schematic diagram of the State Feedback Control (SFC) model and networks impacted in LD. According to the SFC model, laryngeal control is based on an estimate of the current laryngeal state maintained by a comparison between the predicted state and the incoming feedback. State corrections are generated when there is a mismatch between the predicted state and the actual state as conveyed by the feedback signals. Brain regions marked in blue appear to be abnormally impacted in LD. The numbers in purple indicate the time windows in which abnormalities are observed. Abbreviations used: M1 = Primary Motor Cortex, vPMC = Ventral Premotor Cortex, IPL = Inferior Parietal Lobule, S1 = Primary Somatosensory Cortex, A1 = Primary Auditory Cortex, vSMG = Ventral Supramarginal Gyrus. pSTS = Posterior Superior Temporal Sulcus.

The abnormal activation dynamics before and after glottal movement onset together suggest a general impairment in activation of phonation control and initiation of feedforward somatosensory predictions of glottal movement. Prior to glottal movement onset, the observed hypoactivity patterns in patients in the left IFG may indicate abnormalities in activating a phonation control network. Reduced activity in patients in left vPMC, the location where predictions of the estimated laryngeal state are generated, suggests difficulties in sending motor commands to the larynx from M1 and in generating predictions about the somatosensory consequences of these motor commands. Abnormalities in cerebellar activation suggest impaired state prediction. After glottal movement onset, hyperactivity in right ATL, right IFG, bilateral STG and MTG may indicate hyperactivity associated with auditory predictions.

The hyperactivity around voice onset that we see in high-gamma band in auditory cortices may suggest that patients with LD have trouble in setting up auditory predictions in advance of the arrival of auditory feedback at voice onset as well as deficits in early stages of processing auditory feedback.

Midway into pitch perturbation, we also observed high-gamma band hyperactivity in the left posterior parieto-occipital junction, which is involved in multisensory integration. ^75, 76^ During auditory feedback perturbation there is a perceived auditory error that drives a motoric compensation which in turn drives a perceived somatosensory prediction error in the opposite direction. The observed abnormal high-gamma activations can thus be interpreted as abnormal modulation of this somatosensory response.

### Limitations and future directions

An intrinsic limitation of our study was that we could only include data from patients whose voices were functional enough to do the task. This limited the degree of severity that could be studied. Also, the focus of the current study was on the adductor type of LD. Future studies including patients with abductor LD are necessary to investigate whether the findings are specific to adductor LD or reflect a more general LD phenotype. Studies to explain the mechanistic differences between the two spectral bands in this paper should be pursued in the future.

Nevertheless, LD-specific activation abnormalities identified in various brain regions within the speech motor network around various phonation events not only provide temporal-specificity to neuroimaging phenotypes in LD but also may serve as potential therapeutic targets for neuromodulation.

## Abbreviations

CNS: Central Nervous System
LD: Laryngeal Dystonia
MEG: Magnetoencephalography

## Acknowledgements

We would like to thank Anne Findlay, Corby Dale, Rowan Saloner, Mary McPolin, Kim Semien and the study participants.

## Funding

This work was supported by the National Institutes of Health grants R21DC014525 and R01DC013979, grant from the National Spasmodic Dysphonia Association, resident research grant from the American Academy of Otolaryngology - Head and Neck Surgery Foundation to Molly Naunheim, UCSF Discovery Fellowship to Hardik Kothare. These funding sources were not involved in the study design, analysis, interpretation and writing of the article.

## Competing interests

The authors report no competing interests.

## Supplementary material

Supplementary material is available online.

## Notes

### Competing Interest Statement

The authors have declared no competing interest.

