## Supplementary Figures and Tables for "Temporal specificity of abnormal neural oscillations during phonatory events in Laryngeal Dystonia"

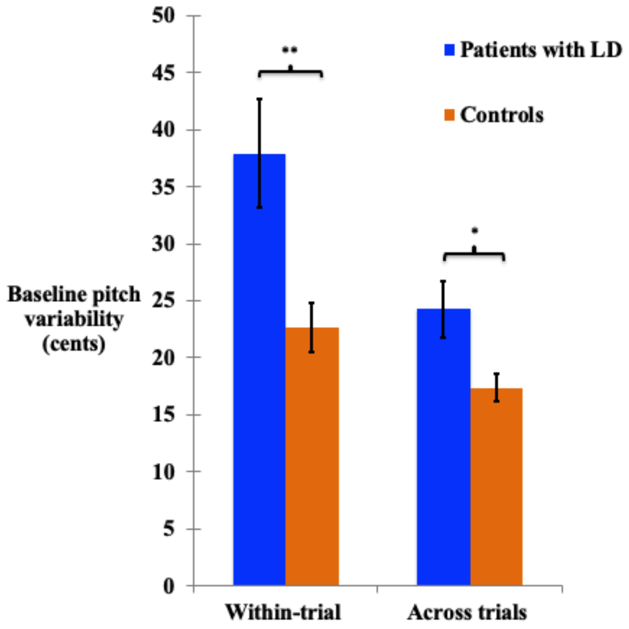


**Supplementary Figure 1: Baseline pitch variability in patients with LD and controls**

Baseline vocal range: Baseline pitch variability (200ms prior to perturbation onset) in patients with LD (n = 15) differs from that in controls (n = 12) both within-trial (two-sample heteroscedastic t-test, t = 2.94, p = 0.0038) and across trials (two-sample heteroscedastic t-test, t = 2.5, p = 0.0202) . Error bars indicate standard error of the mean variability.

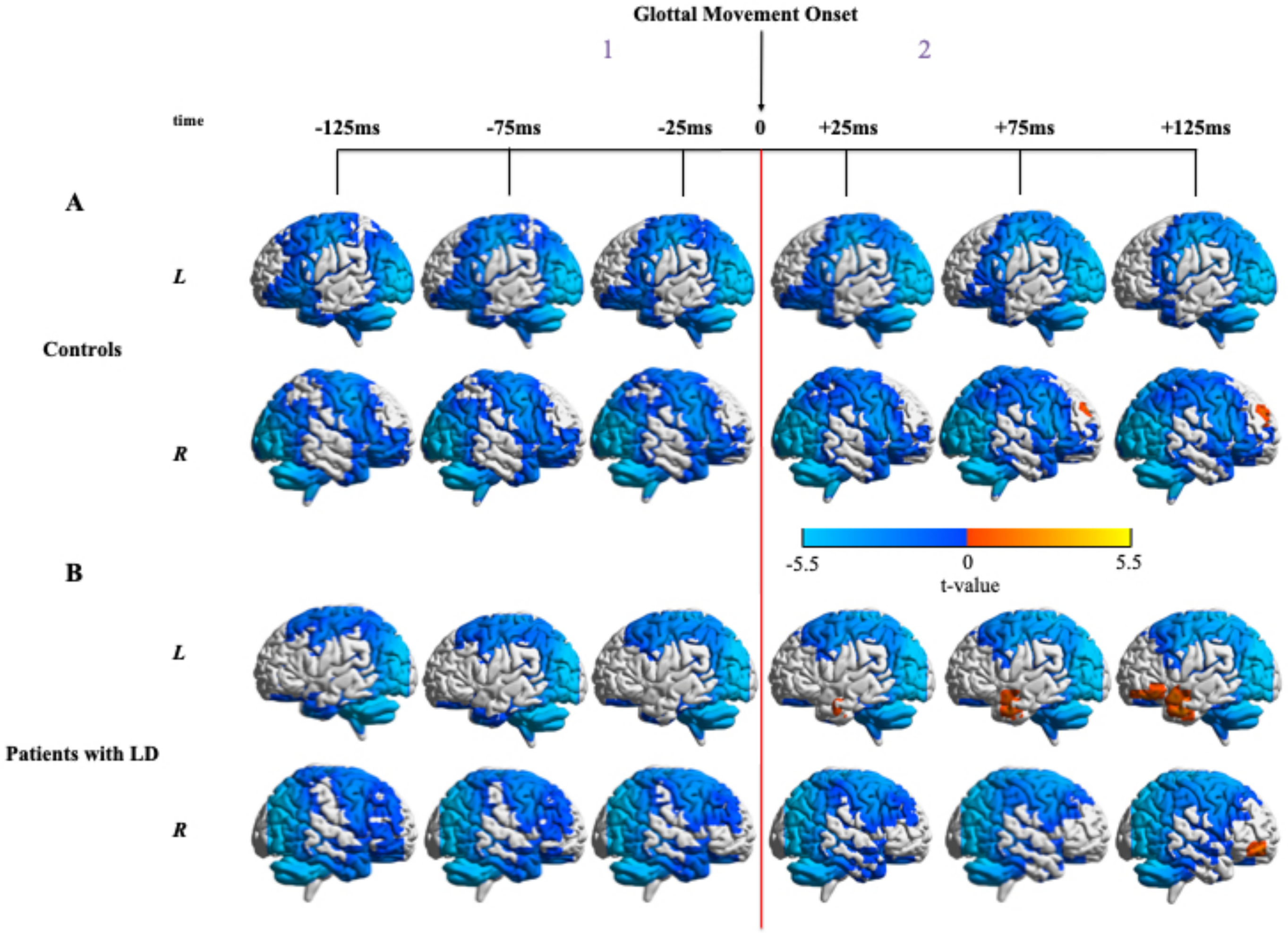


**Supplementary Figure 2: Neural activity in controls and patients with LD in the beta band (12 - 30 Hz) locked to glottal movement onset**

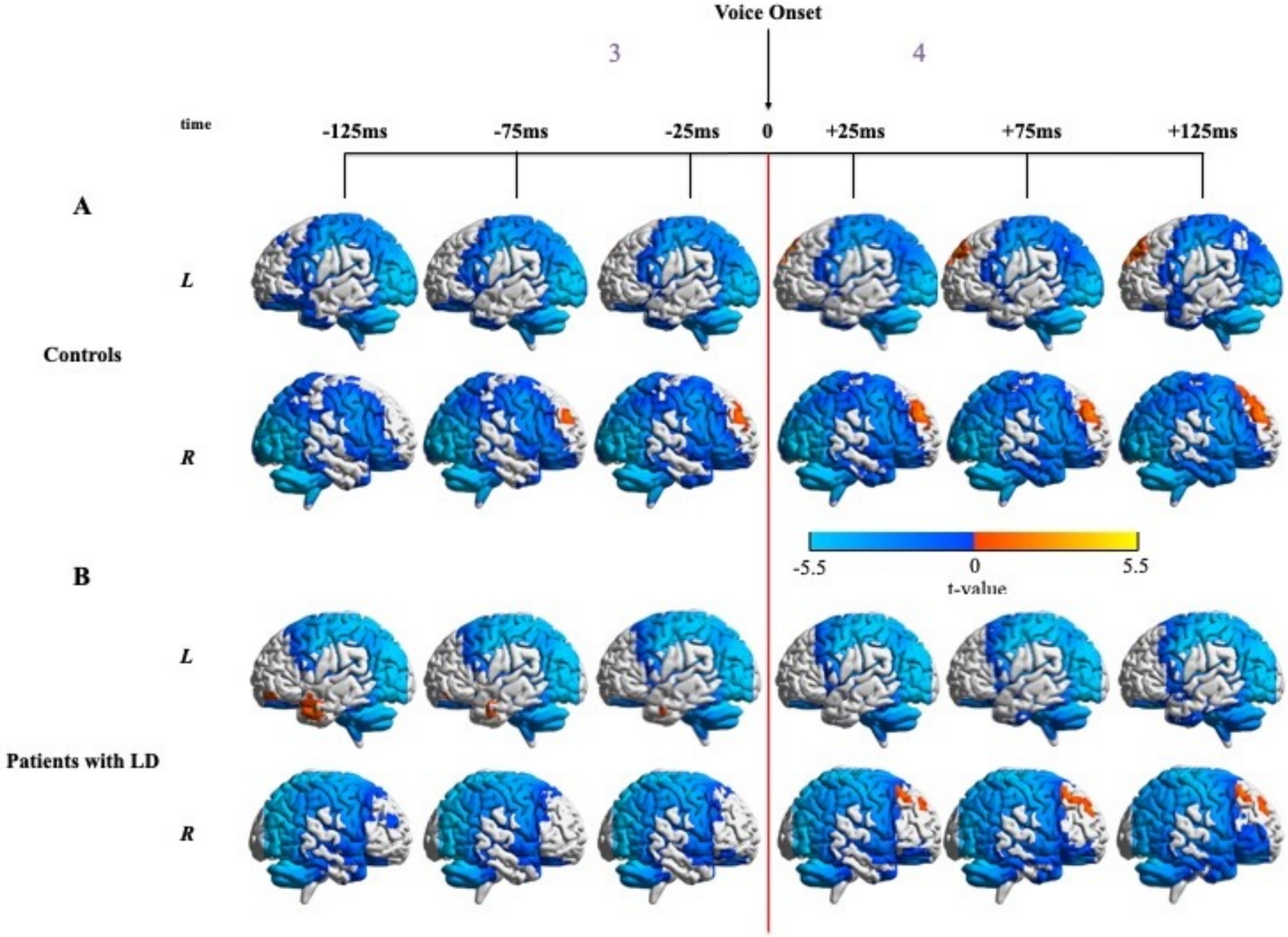


**Supplementary Figure 3: Neural activity in controls and patients with LD in the beta band (12 - 30 Hz) locked to voice onset**

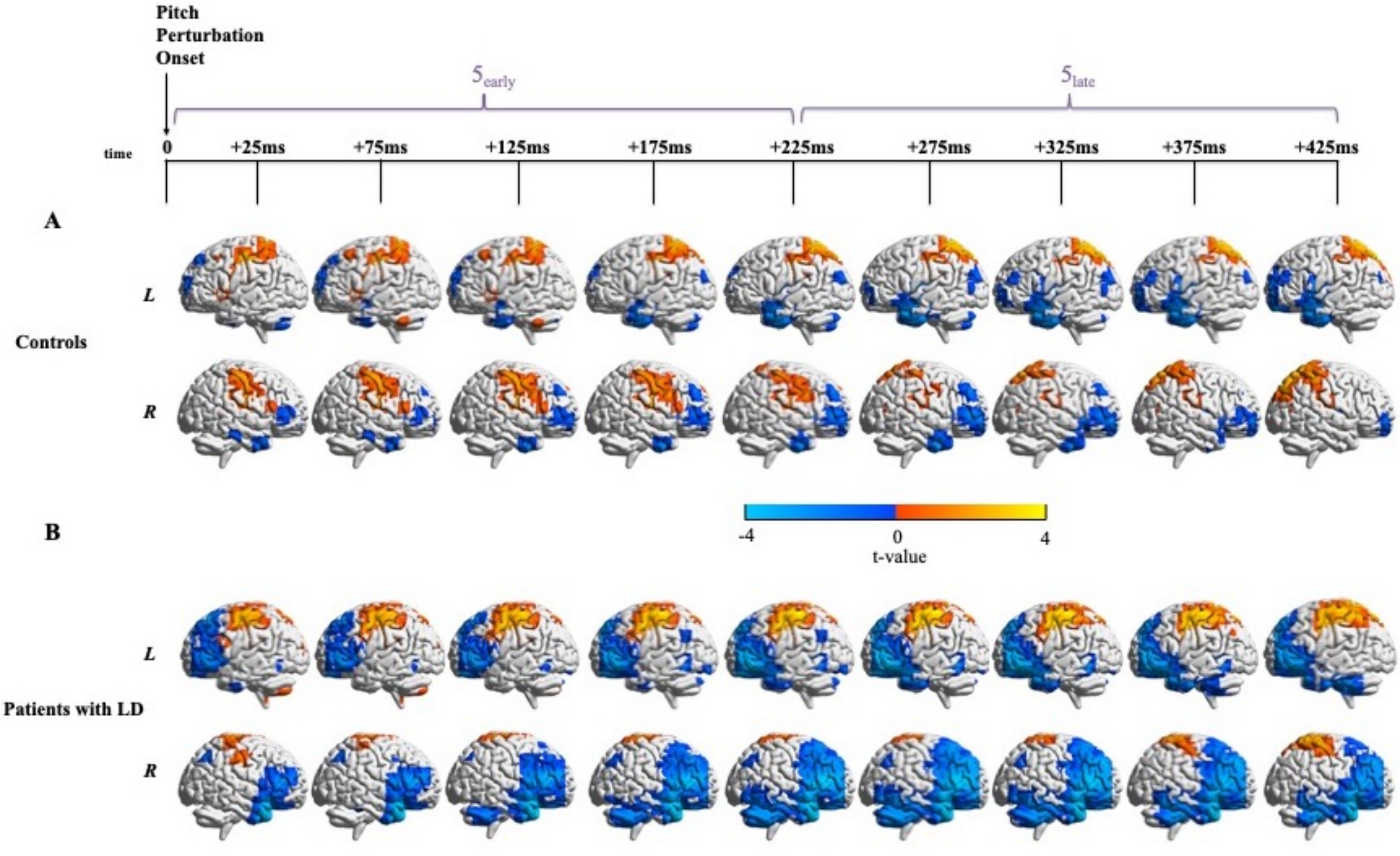


**Supplementary Figure 4: Neural activity in controls and patients with LD in the beta band (12 - 30 Hz) locked to pitch perturbation onset**

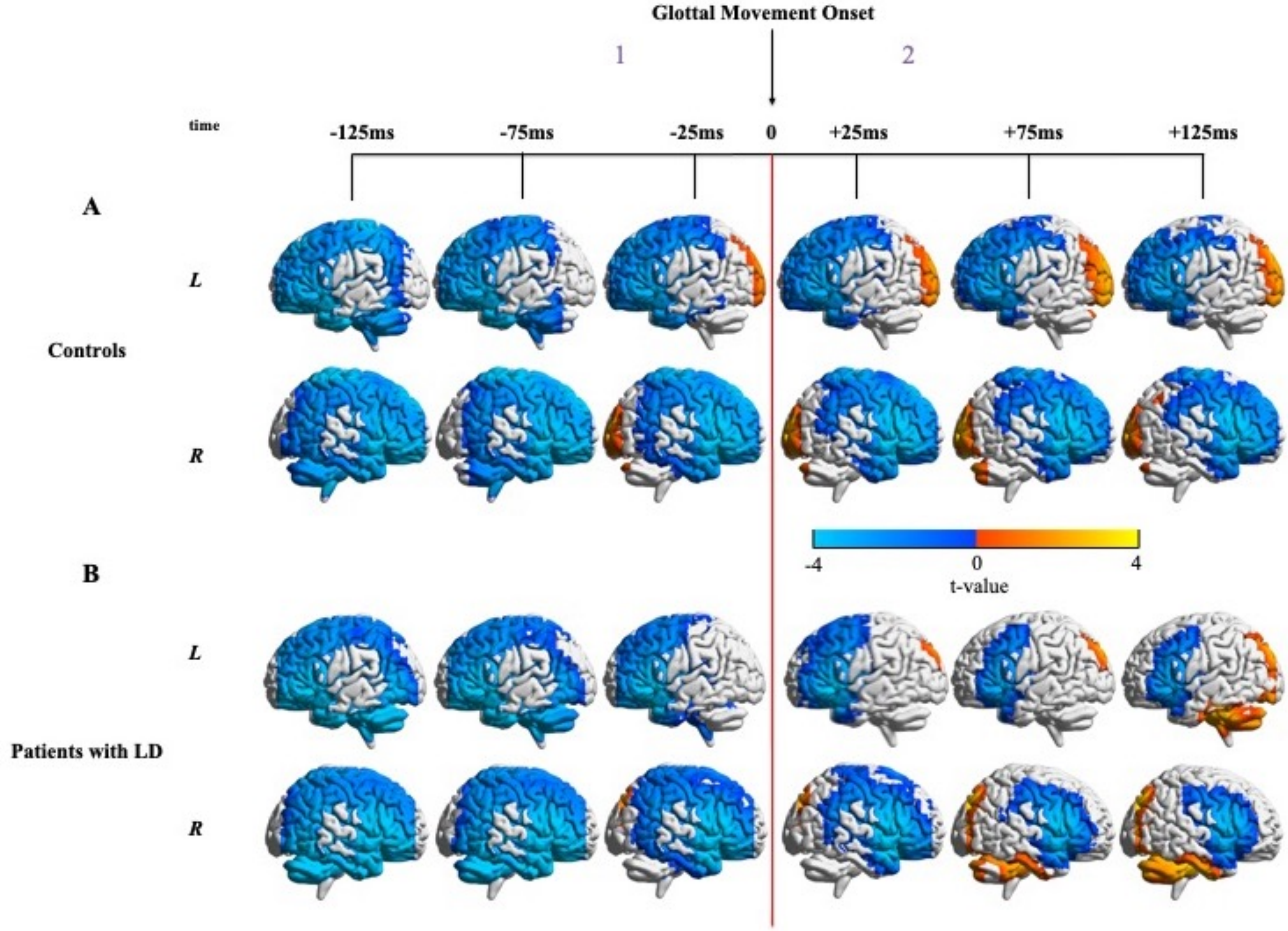


**Supplementary Figure 5: Neural activity in controls and patients with LD in the high gamma band (65-150 Hz) locked to glottal movement onset**

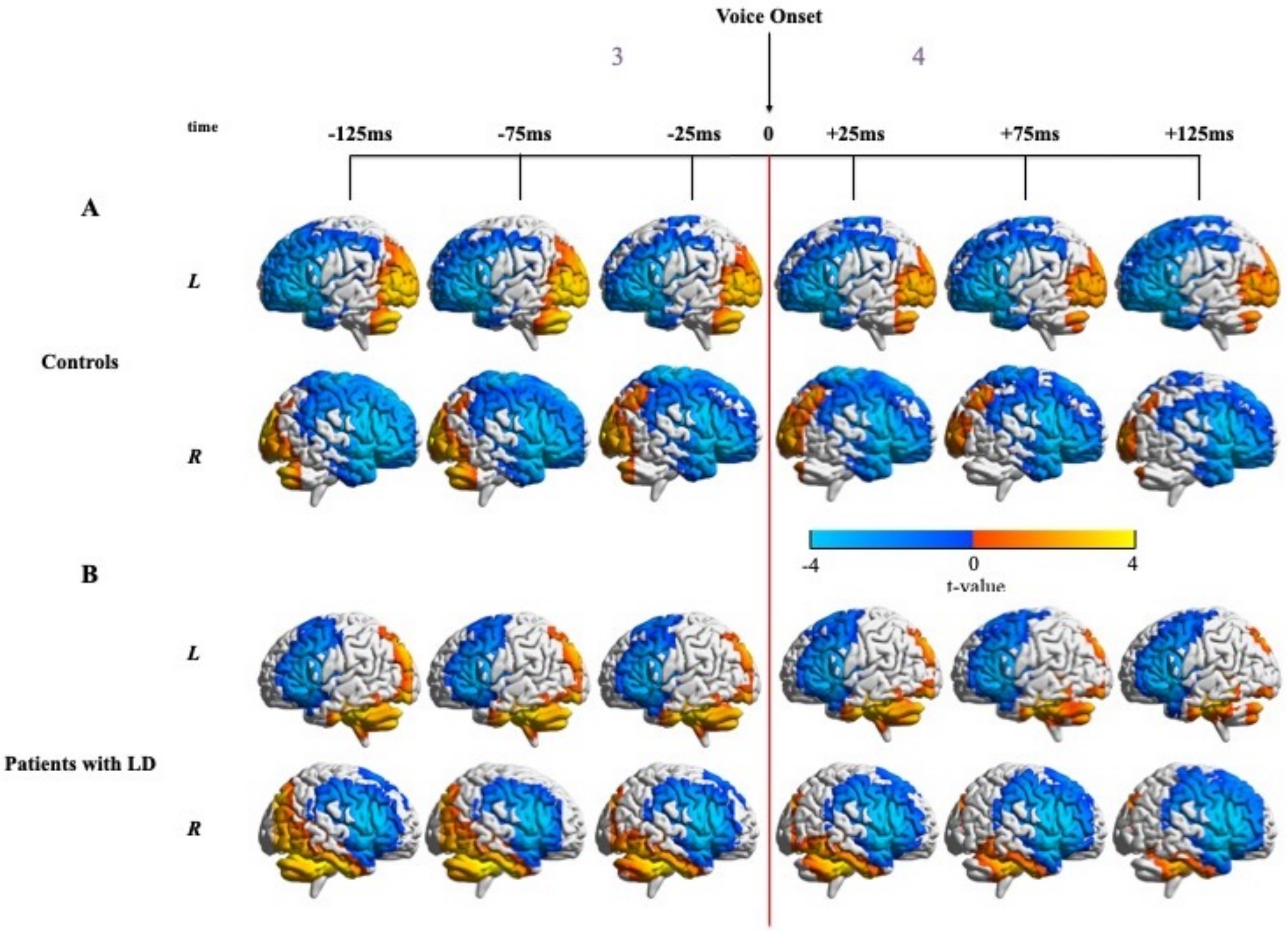


**Supplementary Figure 6: Neural activity in controls and patients with LD in the high gamma band (65-150 Hz) locked to voice onset**

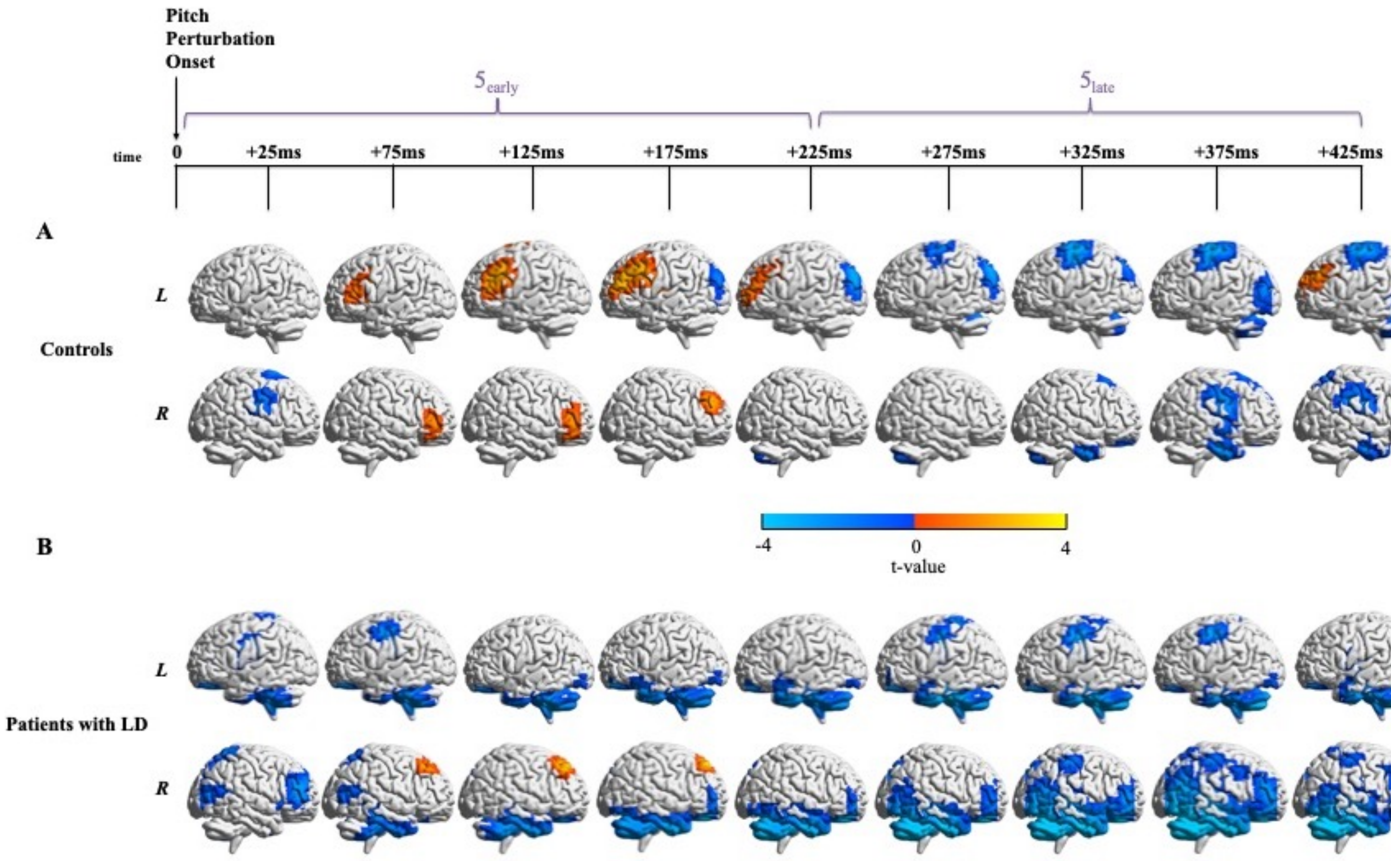


**Supplementary Figure 7: Neural activity in controls and patients with LD in the high gamma band (65-150 Hz) locked to pitch perturbation onset**

**Supplementary Table 1: Meta-analysis of studies of the CNS in patients with Adductor LD**

| Study | Modality | Task | Regions with increase in activity | Regions with decrease in activity | Regions with increase in connectivity (Seed - Target) | Regions with decrease in connectivity (Seed - Target) |
|---|---|---|---|---|---|---|
| Haslinger B, et al. (2005), Neurology; 65(10): 1562-9. | Silent event-related fMRI | Prolonged vowel phonation | | L superior M1 and S1, R inferior M1 and S1, ACC, mesial and L SMA, L and R dPMC, L SFG, R IFG, L MFG, R superior parietal,R occipital, L fusiform gyrus, L parahippocampal gyrus, L hemisphere of the cerebellum | | |
| | | Whispered speech | | L superior M1, R inferior postcentral, L and R inferior parietal, mesial frontal, R STG, R MTG, R fusiform gyrus, R parieto-occipital | | |
| Ali et al. (2006), Journal of Speech, Language, and Hearing Research; Vol. 49 1127-1146 | $H_2{}^{15}O$ PET | Narrative speech | Cerebellum, dorsal and ventral precentral gyrus, anterior insula, ACC, A1, SII, posterior auditory association cortex | Posterior SMG, posterior MTG, dorsal postcentral gyrus, anterior auditory association cortex, PAG, SMA, anterior MTG | | |
| Simonyan & Ludlow (2010), Cerebral Cortex; 20:2749--2759 | fMRI | Symptomatic syllable production | M1, S1, Insula, STG, MTG, Cerebellum, Operculum, Basal Ganglia, Thalamus, | Midbrain | | |
| | | Asymptomatic whimper | M1, S1, Operculum, Insula, MCC, MTG | Insula, Thalamus, Basal Ganglia, Cerebellum, SMA | | |
| | | Coughing | M1, S1, Operculum, anterior insula, STG, MCC, MTG, Midbrain | Cerebellum | | |
| | | Voluntary breathing | M1, S1 | Insula, Midbrain, Cerebellum, SMG | | |
| Kiyuna, A., et al. (2017), J Voice; 31(3): p. 379 e1-379 e11. | fMRI | Reading five-digit numbers | L MTG, L thalamus, L and R precentral gyrus, L and R postcentral gyrus, R insula, R Cerebellum VIII and IX, R putamen, L Cerebellum I-IV, R SMA | L Cerebellum Crus 1 and 2, R Cerebellum Crus 1, L STG, L Cerebellum VI | | |
| | | Resting state | | | L Thalamus - L Caudate; R precentral gyrus - L ITG, L temporal pole, L MTG; L postcentral gyrus - R frontal pole; L inferior operculum - R precentral and postcentral gyri; Cerebellum (vermis I, II) - R lateral occipital cortex, R superior parietal lobule; R Cerebellum (IX) - R precentral and postcentral gyri | L insula - R angular gyrus, R lateral occipital cortex; R thalamus - L MFG, L IFG, pars triangularis; L precuneus - L and R lingual gyrus; R precentral gyrus - R occipital pole |
| Khosravani S, et al. (2019), Clinical Neurophysiology; 130(6): 1033-40. | EEG | Vowel vocalisation | L somatosensory-premotor cortices (late vocalisation, gamma band) | L motor cortex (early vocalisation, alpha band) | | |
| Daliri A, et al. (2020), Journal of Speech, Language, and Hearing Research; 63(2): 421-32. | fMRI | Normal sentence production and Sentence production under masking noise | L ventral sensorimotor cortex, L anterior planum temporale, L posterior STG / planum temporale | | | |
| | | Resting state | | | L mid-Rolandic cortex - L Heschl's gyrus, L posterior STG, R Heschl's gyrus; L ventral Rolandic cortex - L posterior STG; R mid-Rolandic cortex - L posterior STG | |

Abbreviations used: fMRI = Functional Magnetic Resonance Imaging, PET = Positron Emission Tomography, EEG = Electroencephalography, L = left, R = right, M1 = primary motor cortex, S1 = primary somatosensory cortex, ACC = anterior cingulate cortex, SMA = supplementary motor area, dPMC = dorsal premotor cortex, SFG = superior frontal gyrus, IFG = inferior frontal gyrus, MFG = middle frontal gyrus, STG = superior temporal gyrus, MTG = middle temporal gyrus, A1 = primary auditory cortex, SII = secondary somatosensory cortex, SMG = supramarginal gyrus, PAG = periaqueductal grey, MCC = middle cingulate cortex, ITG = inferior temporal gyrus.

**Supplementary Table 2: Peak voxels with significant beta-band activity differences with respect to glottal movement onset**

| Hemisphere | Peak | MNI Coordinates | Anatomical Labels | Time with respect to glottal movement onset |
|---|---|---|---|---|
| Left | 1 | [-41.2 -78.1 32.2] | Angular Gyrus (Brodmann Area 39) | -125 to +125ms |
| | 2 | [-40.4 -81.9 28.3] | Superior Occipital Gyrus (Brodmann Area 19) | -125 to +125ms |
| | 3 | [-41.9 -70.3 25.9] | Middle Temporal Gyrus (Brodmann Area 39) | -125ms |
| | 4 | [-23.3 -35.8 -25.7] | Cerebellar Anterior Lobe | -125ms |
| | 5 | [-59.0 -3.8 22.0] | Precentral Gyrus (Brodmann Area 4) | -125 to +25ms |
| | 6 | [-59.0 2.5 22.0] | Precentral Gyrus (Brodmann Area 6) | -125 to +75ms |
| | 7 | [-63.7 -29.0 27.5] | Inferior Parietal Lobule (Brodmann Area 40) | -125 to -25ms |
| | 8 | [-55.2 19.6 -18.4] | Superior Temporal Gyrus (Brodmann Area 38) | -125 to +125ms |
| | 9 | [-55.2 28.1 -8.9] | Inferior Frontal Gyrus (Brodmann Area 47) | -125 to +125ms |
| | 10 | [-55.9 4.1 20.4] | Inferior Frontal Gyrus (Brodmann Area 6) | -125 to +75ms |
| | 11 | [-48.2 -76.4 26.7] | Middle Temporal Gyrus (Brodmann Area 39) | -75 to +125ms |
| | 12 | [-58.3 3.3 15.6] | Inferior Frontal Gyrus (Brodmann Area 44) | +75 to +125ms |
| | 13 | [-51.3 21.9 8.5] | Inferior Frontal Gyrus (Brodmann Area 45) | +75ms |
| | 14 | [-47.4 43.5 -17.6] | Inferior Frontal Gyrus (Brodmann Area 47) | +125ms |
| | 15 | [-14.8 -48.6 -57.2] | Cerebellar Tonsil | +125ms |
| | 16 | [-59.0 5.7 12.5] | Precentral Gyrus (Brodmann Area 44) | +125ms |
| | 17 | [-52.0 12.0 -24.7] | Superior Temporal Gyrus (Brodmann Area 38) | +125ms |
| | 18 | [-52.0 9.5 -30.2] | Middle Temporal Gyrus (Brodmann Area 38) | +125ms |
| | 19 | [-30.3 17.3 60.7] | Middle Frontal Gyrus (Brodmann Area 6) | +125ms |
| Right | 1 | [41.9 -69.1 34.6] | Precuneus (Brodmann Area 39) | -125 to +25ms |
| | 2 | [41.9 -69.1 39.3] | Inferior Parietal Lobule (Brodmann Area 39) | -125 to +125ms |
| | 3 | [49.0 -72.2 37.0] | Angular Gyrus (Brodmann Area 39) | -125 to +25ms |
| | 4 | [60.1 -27.9 -27.1] | Inferior Temporal Gyrus (Brodmann Area 20) | -125ms |
| | 5 | [24.5 25.2 62.3] | Superior Frontal Gyrus (Brodmann Area 8) | -25 to +125ms |
| | 6 | [16.6 -77.8 34.6] | Precuneus (Brodmann Area 19) | +75ms |
| | 7 | [8.0 -83.0 47.0] | Precuneus (Brodmann Area 7) | +75 to +125ms |
| | 8 | [59.3 23.6 -1.0] | Inferior Frontal Gyrus (Brodmann Area 45) | +75 to +125ms |
| | 9 | [48.2 52.1 -8.9] | Middle Frontal Gyrus (Brodmann Area 10) | +75 to +125ms |
| | 10 | [31.6 -84.1 -14.4] | Middle Occipital Gyrus (Brodmann Area 19) | +75 to +125ms |
| | 11 | [23.7 -77.0 -24.7] | Cerebellar Posterior Lobe | +75 to +125ms |
| | 12 | [22.1 -75.4 -9.7] | Lingual Gyrus | +75 to +125ms |
| | 13 | [49.8 -58.0 -16.8] | Fusiform Gyrus | +125ms |
| | 14 | [39.5 -53.2 44.9] | Inferior Parietal Lobule | +125ms |

**Supplementary Table 3: Peak voxels with significant high-gamma-band activity differences with respect to glottal movement onset**

| Hemisphere | Peak | MNI Coordinates | Anatomical Labels | Time with respect to glottal movement onset |
|---|---|---|---|---|
| Left | 1 | [-64.0 -27.0 39.0] | Postcentral Gyrus (Brodmann Area 40) | +25 to +125ms |
| | 2 | [-65.3 -25.5 -8.4] | Middle Temporal Gyrus (Brodmann Area 21) | +125ms |
| | 3 | [-61.4 5.6 -2.9] | Superior Temporal Gyrus (Brodmann Area 22) | +125ms |
| Right | 1 | [46.6 -74.2 -57] | Cerebellar Posterior Lobe | -125 to -75ms |
| | 2 | [48.0 37.0 -9.0] | Inferior Frontal Gyrus (Brodmann Area 47) | +75 to +125ms |
| | 3 | [46.6 4.4 -40.6] | Middle Temporal Gyrus (Brodmann Area 21) | +75 to +125ms |
| | 4 | [62.1 -26.9 8.2] | Superior Temporal Gyrus (Brodmann Area 41) | +125ms |

**Supplementary Table 4: Peak voxels with significant beta-band activity differences with respect to voice onset**

| Hemisphere | Peak | MNI Coordinates | Anatomical Labels | Time with respect to voice onset |
|---|---|---|---|---|
| Left | 1 | [-54.4 14.2 -27.1] | Superior Temporal Gyrus (Brodmann Area 38) | -125 to -25ms |
| | 2 | [-54.4 9.5 -32.6] | Middle Temporal Gyrus (Brodmann Area 38) | -125 to -75ms |
| | 3 | [-33.4 42.0 -17.6] | Middle Frontal Gyrus (Brodmann Area 47) | -125ms |
| | 4 | [-55.2 20.4 14.0] | Inferior Frontal Gyrus (Brodmann Area 45) | -125 to -75ms |
| | 5 | [-31.8 11.1 60.7] | Middle Frontal Gyrus (Brodmann Area 6) | -125 to +25ms |
| | 6 | [-14.8 -13.7 68.6] | Superior Frontal Gyrus (Brodmann Area 6) | -125 to -75ms |
| | 7 | [-38.8 -27.5 45.7] | Postcentral Gyrus (Brodmann Area 2) | -125 to -75ms |
| | 8 | [-39.6 -76.5 38.6] | Precuneus (Brodmann Area 19) | -125 to -75ms |
| | 9 | [-39.6 -83.5 27.5] | Superior Occipital Gyrus (Brodmann Area 19) | -125 to -75ms |
| | 10 | [-42.7 -79.5 30.6] | Angular Gyrus (Brodmann Area 39) | -125 to -75ms |
| | 11 | [-31.8 -27.5 69.4] | Precentral Gyrus (Brodmann Area 4) | -25 to +125ms |
| | 12 | [-52.8 -26.7 58.3] | Postcentral Gyrus (Brodmann Area 1) | -25 to +125ms |
| | 13 | [-31.1 -85.0 -11.2] | Inferior Occipital Gyrus (Brodmann Area 18) | -25 to +125ms |
| | 14 | [-15.5 -58.7 -57.9] | Cerebellar Tonsil | -25 to +125ms |
| Right | 1 | [22.9 -66.7 62.3] | Superior Parietal Lobule (Brodmann Area 7) | -125 to -75ms |
| | 2 | [22.1 -62.0 61.5] | Superior Parietal Lobule (Brodmann Area 7) | -125 to -75ms |
| | 3 | [34.0 -35.8 69.4] | Postcentral Gyrus (Brodmann Area 1) | -125 to +125ms |
| | 4 | [25.3 -84.9 -16.8] | Middle Occipital Gyrus (Brodmann Area 19) | -125 to +125ms |
| | 5 | [56.9 -59.6 -17.6] | Inferior Temporal Gyrus (Brodmann Area 37) | -25 to +125ms |
| | 6 | [47.4 -50.9 -50.0] | Cerebellar Tonsil | -25 to +125ms |
| | 7 | [34.0 -28.7 67.0] | Precentral Gyrus (Brodmann Area 4) | +25 to +125ms |
| | 8 | [6.3 -3.3 67.0] | Superior Frontal Gyrus (Brodmann Area 6) | +25 to +75ms |
| | 9 | [53.7 20.4 13.3] | Inferior Frontal Gyrus (Brodmann Area 44) | +75 to +125ms |
| | 10 | [41.1 22.0 37.0] | Middle Frontal Gyrus (Brodmann Area 8) | +75 to +125ms |
| | 11 | [6.3 3.8 69.4] | Superior Frontal Gyrus (Brodmann Area 6) | +125ms |

**Supplementary Table 5: Peak voxels with significant high-gamma-band activity differences with respect to voice onset**

| Hemisphere | Peak | MNI Coordinates | Anatomical Labels | Time with respect to voice onset |
|---|---|---|---|---|
| Left | 1 | [-8.0 53.0 39.0] | Superior Frontal Gyrus (Brodmann Area 9) | -125ms |
| | 2 | [-64.0 -19.0 23.0] | Postcentral Gyrus (Brodmann Area 1) | -125 to -25ms |
| | 3 | [-16.0 61.0 -17.0] | Superior Frontal Gyrus (Brodmann Area 11) | -125 to +125ms |
| | 4 | [-56.0 -11.0 -17.0] | Middle Temporal Gyrus (Brodmann Area 21) | -125 to +125ms |
| Right | 1 | [41.3 6.5 -42.2] | Inferior Temporal Gyrus (Brodmann Area 20) | -125 to +25ms |
| | 2 | [48.0 -19.0 -33.0] | Inferior Temporal Gyrus (Brodmann Area 20) | -125 to +25ms |
| | 3 | [64.0 -27.5 8.2] | Superior Temporal Gyrus (Brodmann Area 41) | -125 to +125ms |
| | 4 | [8.0 -67.0 39.0] | Precuneus (Brodmann Area 7) | -125 to -75ms |
| | 5 | [46.8 -68.5 45.2] | Inferior Parietal Lobule (Brodmann Area 39) | +75ms |

**Supplementary Table 6: Peak voxels with significant beta-band activity differences with respect to pitch perturbation onset**

| Hemisphere | Peak | MNI Coordinates | Anatomical Labels | Time with respect to voice onset |
|---|---|---|---|---|
| Left | 1 | [-39.6 -75.0 -50.8] | Cerebellar Inferior Semi-Lunar Lobule | +25 to +225ms |
| | 2 | [-48.2 20.4 4.6] | Inferior Frontal Gyrus (Brodmann Area 45) | +25 to +175ms |
| | 3 | [-38.8 23.5 44.1] | Middle Frontal Gyrus (Brodmann Area 8) | +25ms |
| | 4 | [-25.6 -59.0 44.9] | Superior Parietal Lobule (Brodmann Area 7) | +175 to +375ms |
| | 5 | [-52.8 27.0 -11.2] | Inferior Frontal Gyrus (Brodmann Area 47) | +225ms |
| | 6 | [-25.6 -51.9 43.3] | Precuneus (Brodmann Area 7) | +225 to +375ms |
| | 7 | [-46.6 34.9 -11.2] | Inferior Frontal Gyrus (Brodmann Area 47) | +275ms |
| Right | 1 | [48.0 21.0 -25.0] | Superior Temporal Gyrus (Brodmann Area 38) | +25 to +425ms |
| | 2 | [62.4 26.0 10.1] | Inferior Frontal Gyrus (Brodmann Area 9) | +25 to +425ms |
| | 3 | [54.5 20.4 14.8] | Inferior Frontal Gyrus (Brodmann Area 44) | +25 to +325ms |
| | 4 | [49.8 -3.3 33.8] | Precentral Gyrus (Brodmann Area 6) | +75 to +275ms |
| | 5 | [49.8 3.8 38.6] | Middle Frontal Gyrus (Brodmann Area 6) | +125 to +275ms |
| | 6 | [55.3 3.8 31.4] | Inferior Frontal Gyrus (Brodmann Area 6) | +125 to +275ms |
| | 7 | [55.3 26.0 14.0] | Inferior Frontal Gyrus (Brodmann Area 9) | +125 to +325ms |
| | 8 | [55.3 -67.5 -41.3] | Cerebellar Posterior Lobe | +175 to +325ms |
| | 9 | [18.2 52.9 38.6] | Superior Frontal Gyrus (Brodmann Area 9) | +225 to +325ms |
| | 10 | [15.0 62.4 7.7] | Superior Frontal Gyrus (Brodmann Area 10) | +375 to +425ms |
| | 11 | [55.3 -52.5 -43.7] | Cerebellar Tonsil | +375 to +425ms |
| | 12 | [45.8 -66.7 45.7] | Inferior Parietal Lobule (Brodmann Area 39) | +375 to +425ms |
| | 13 | [63.2 -12.1 -16.0] | Middle Temporal Gyrus (Brodmann Area 21) | +425ms |

**Supplementary Table 7: Peak voxels with significant high-gamma-band activity differences with respect to pitch perturbation onset**

| Hemisphere | Peak | MNI Coordinates | Anatomical Labels | Time with respect to pitch perturbation onset |
|---|---|---|---|---|
| Left | 1 | [-32.0 29.0 7.0] | Inferior Frontal Gyrus (Brodmann Area 45) | +25 to +125ms |
| | 2 | [10/9 35.1 -26.5] | Rectal Gyrus (Brodmann Area 11) | +25 to +425ms |
| | 3 | [-38.1 -42.0 -54.0] | Cerebellar Tonsil | +25 to +425ms |
| | 4 | [-31.1 -81.9 38.1] | Angular Gyrus (Brodmann Area 39) | +175 to +275ms |
| | 5 | [-48.0 -20.7 -34.3] | Inferior Temporal Gyrus (Brodmann Area 20) | +225 to +325ms |
| Right | 1 | [52.3 37.2 17.6] | Middle Frontal Gyrus (Brodmann Area 46) | +25 to +225ms |
| | 2 | [14.6 29.3 -25.0] | Orbital Gyrus (Brodmann Area 47) | +125 to +425ms |
| | 3 | [56.0 -27.5 -24.9] | Inferior Temporal Gyrus (Brodmann Area 20) | +125 to +425ms |
| | 4 | [48.0 45.0 23.0] | Middle Frontal Gyrus (Brodmann Area 46) | +125 to +225ms |
| | 5 | [56.3 -74.0 -33.0] | Cerebellar Posterior Lobe | +175 to +425ms |
| | 6 | [8.0 -91.0 31.0] | Cuneus (Brodmann Area 19) | +175 to +275ms |
| | 7 | [46.0 -84.3 -16.2] | Inferior Occipital Gyrus (Brodmann Area 18) | +225 to +425ms |
| | 8 | [24.0 33.3 54.6] | Superior Frontal Gyrus (Brodmann Area 8) | +325ms |
