## Supplementary Tables for "Temporal specificity of abnormal neural oscillations during phonatory events in Laryngeal Dystonia"

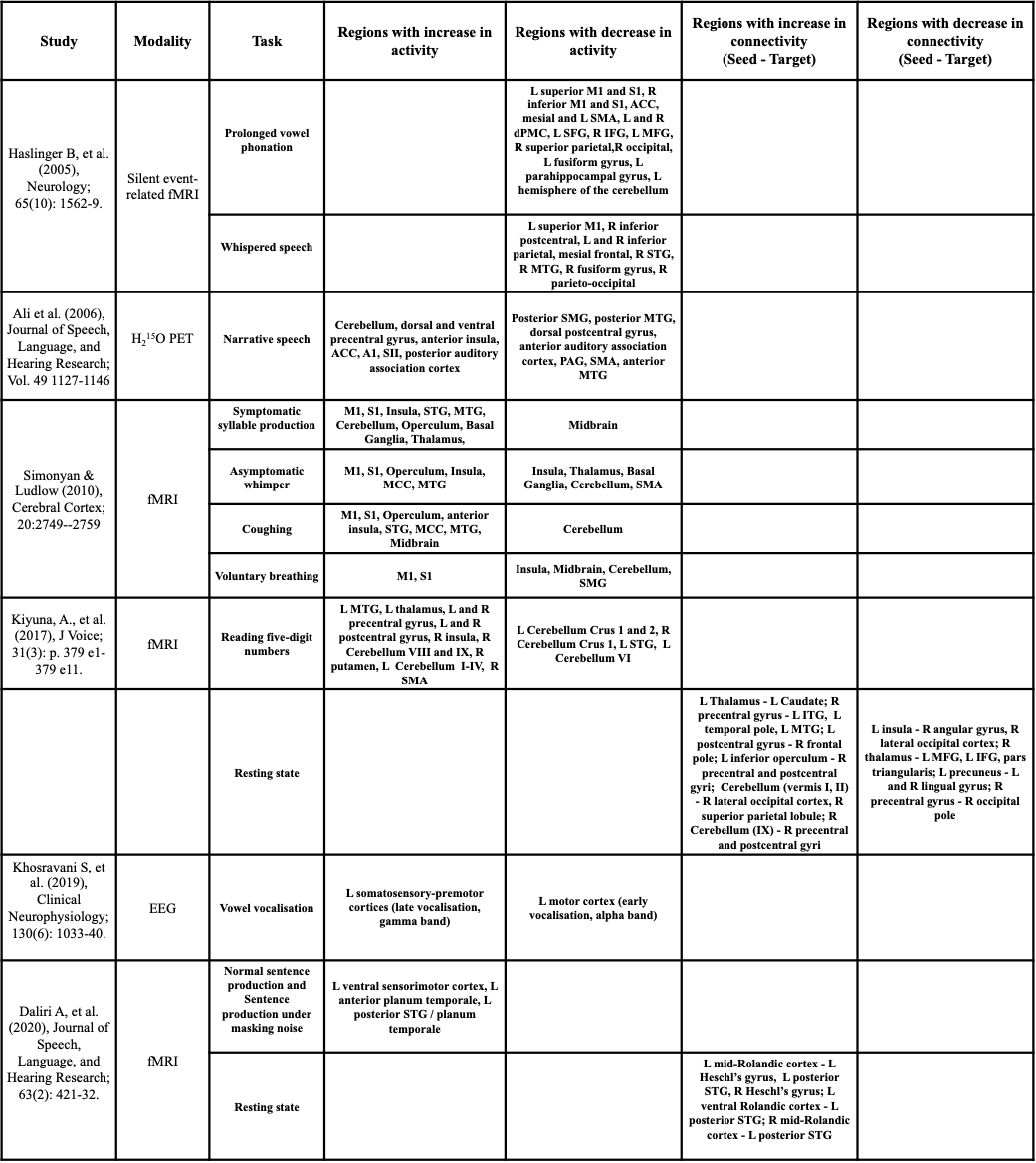

**Supplementary Table 1: Meta-analysis of studies of the CNS in patients with Adductor LD**

Abbreviations used: fMRI = Functional Magnetic Resonance Imaging, PET = Positron Emission Tomography, EEG = Electroencephalography, L = left, R = right, M1 = primary motor cortex, S1 = primary somatosensory cortex, ACC = anterior cingulate cortex, SMA = supplementary motor area, dPMC = dorsal premotor cortex, SFG = superior frontal gyrus, IFG = inferior frontal gyrus, MFG = middle frontal gyrus, STG = superior temporal gyrus, MTG = middle temporal gyrus, A1 = primary auditory cortex, SII = secondary somatosensory cortex, SMG = supramarginal gyrus, PAG = periaqueductal grey, MCC = middle cingulate cortex, ITG = inferior temporal gyrus.

**Supplementary Table 2: Number of participants in each analysis**

| **Analysis** | **Controls** | **Patients** |
| --- | --- | --- |
| Phonatory onset interval | 11 | 15 |
| Beta band and high gamma band MEG activity around glottal movement onset | 11 | 15 |
| Beta band and high gamma band MEG activity around voice onset | 11 | 15 |
| Behavioural response to pitch perturbation onset | 12 | 17 |
| Correlation between mean compensation to pitch perturbation and laryngeal diadochokinesis rate^a^ | 11 | 12 |
| Beta band MEG activity after pitch perturbation onset | 11 | 15 |
| High gamma band MEG activity after pitch perturbation onset^b^ | 16 | 15 |

^a^The number of participants is lower for this analysis because only those participants who underwent clinical voice evaluation and had good vocal response data could be included.

^b^Five controls were added to this analysis to improve signal-to-noise ratio.
